# Cyb5r3 links FoxO1-dependent mitochondrial dysfunction with β-cell failure

**DOI:** 10.1101/774364

**Authors:** Jason Fan, Wen Du, Ja Young Kim-Muller, Jinsook Son, Taiyi Kuo, Delfina Larrea, Christian Garcia, Takumi Kitamoto, Michael J. Kraakman, Edward Owusu-Ansah, Vincenzo Cirulli, Domenico Accili

**Affiliations:** Naomi Berrie Diabetes Center and Departments of Medicine, Columbia University, New York, New York 10032, USA; Genetics and Development, Columbia University, New York, New York 10032, USA; Physiology and Cellular Biophysics, Columbia University, New York, New York 10032, USA; Department of Medicine, UW-Diabetes Institute, Institute for Stem Cell and Regenerative Medicine, University of Washington, Seattle, WA, USA

## Abstract

**Objective:** Diabetes is characterized by pancreatic β-cell dedifferentiation. Dedifferentiating β-cells inappropriately metabolize lipids over carbohydrates and exhibit impaired mitochondrial oxidative phosphorylation. However, the mechanism linking the β-cell’s response to an adverse metabolic environment with impaired mitochondrial function remains unclear.

**Methods:** Here we report that the oxidoreductase cytochrome b5 reductase 3 (Cyb5r3) links FoxO1 signaling to β-cell stimulus/secretion coupling by regulating mitochondrial function, reactive oxygen species generation, and NAD/NADH ratios.

**Results:** Expression of Cyb5r3 is decreased in FoxO1-deficient β-cells. Mice with β-cell-specific deletion of Cyb5r3 have impaired insulin secretion resulting in glucose intolerance and diet-induced hyperglycemia. Cyb5r3-deficient β-cells have a blunted respiratory response to glucose and display extensive mitochondrial and secretory granule abnormalities, consistent with altered differentiation. Moreover, FoxO1 is unable to maintain expression of key differentiation markers in Cyb5r3-deficient β-cells, suggesting that Cyb5r3 is required for FoxO1-dependent lineage stability.

**Conclusions:** The findings highlight a pathway linking FoxO1 to mitochondrial dysfunction that can mediate β-cell failure.

## Introduction

A hallmark of diabetes is β-cell failure, a state of chronically increased demand for insulin that exceeds the β-cell’s secretory capacity and leads to a gradual deterioration of β-cell mass and function [1; 2]. This decline manifests as a mistimed and diminished response to nutrients and, combined with peripheral insulin resistance, gives rise to hyperglycemia [3]. Although incretin-based therapies improve β-cell performance and Glp1 agonists show increased durability compared to other treatments [4], they are insufficient to halt disease progression [5; 6].

There are several pathways to β-cell failure [2]. Our work focuses on β-cell dedifferentiation as a mechanism of β-cell dysfunction [7–13]. We and others have described a signaling mechanism by which excessive or altered nutrient flux, in combination with an adverse hormonal or inflammatory milieu, results in a stress response orchestrated by (but not limited to) the transcription factor FoxO1 [14; 15]. The failure of this homeostatic mechanism affects mitochondrial function [16]. We have proposed that dedifferentiation is the end result of altered mitochondrial substrate utilization, or “metabolic inflexibility” [17]. The implication of this model is that therapeutic reversal of mitochondrial dysfunction may prevent dedifferentiation or even promote “re-differentiation” of β-cells. However, the effectors of mitochondrial dysfunction in this model remain unclear. In previous work, we used a combination of marker analysis and RNA profiling to compile a list of potential effector genes of β-cell failure [18]. Although several interesting candidates emerged from that work, functional evidence for their role in diabetes progression is lacking.

Among candidates identified in the RNA profiling of dedifferentiating β-cells is Cyb5r3. This gene encodes a flavoprotein with membrane-bound and soluble forms, the latter of which is restricted to erythrocytes [19]. Mutations of this form lead to recessive congenital (Type I) methemoglobinemia, while mutations in the membrane-bound form can cause severe neurological disease (Type II) [20]. Membrane-bound Cyb5r3 participates in mitochondrial electron transport chain (ETC) activity, antioxidant reduction, fatty acid desaturation, and cholesterol biosynthesis [19; 21]. Notably, knockout of the related isoform Cyb5r4 causes early-onset diabetes in mice independent of peripheral insulin sensitivity [22]. There are five genes encoding Cyb5r isoforms (R1-4 and RL). In humans, Cyb5r3 is the main isoform expressed in β-cells [23], and its expression increases in iPS-derived β-cells as they undergo terminal differentiation [24]. A SNP in the *CYB5R3* promoter (43049014 G/T) is associated with fasting glucose (*P*= 2.99 × 10-4), consistent with a potential metabolic role [25]. In addition, we have identified a conserved super-enhancer in mouse β-cells and human islets at this locus [26]. Based on this rationale, we tested the hypothesis that Cyb5r3 is a critical FoxO1 target, linking FoxO1 signaling to β-cell mitochondrial ETC function, affecting the generation of reactive oxygen species, NAD/NADH ratios, and stimulus/secretion coupling. Using genetic manipulation of Cyb5r3, we show that the decreased expression of this protein observed in FoxO1-deficient β-cells recapitulates key aspects of FoxO1-dependent β-cell failure.

## Results

### Regulation of β-cell Cyb5r3 gene expression in diabetic mice

During the progression of β-cell failure, loss of FoxO1 expression marks the transition to β-cell dedifferentiation [7; 27-31]. Thus, we reasoned that by identifying FoxO1 target genes in dedifferentiating β-cells, we can identify effectors of β-cell failure. RNAseq experiments indicated that *Cyb5r3* levels are decreased in FoxO knockout β-cells [18]. Nonetheless, we didn’t know whether *Cyb5r3* is a *bona fide* FoxO1 target that mediates β-cell function. To answer this question, we investigated the relationship between FoxO1 activity and Cyb5r3 levels, and determined the occupancy of the *Cyb5r3* promoter by FoxO1.

To validate the previous RNAseq findings of decreased *Cyb5r3*, we purified β-cells from WT and β-cell-specific FoxO knockout mice using a previously described procedure to enrich for dedifferentiating β-cells based on elevated Aldh1A3 activity [18] and measured RNA levels. We found a ∼90% decrease in Cyb5r3 mRNA (Fig. 1A). Moreover, immunohistochemistry of islets from β-cell-specific FoxO1 knockouts [7] also showed decreased intensity of Cyb5r3 (Fig. 1B).

**Figure 1.**
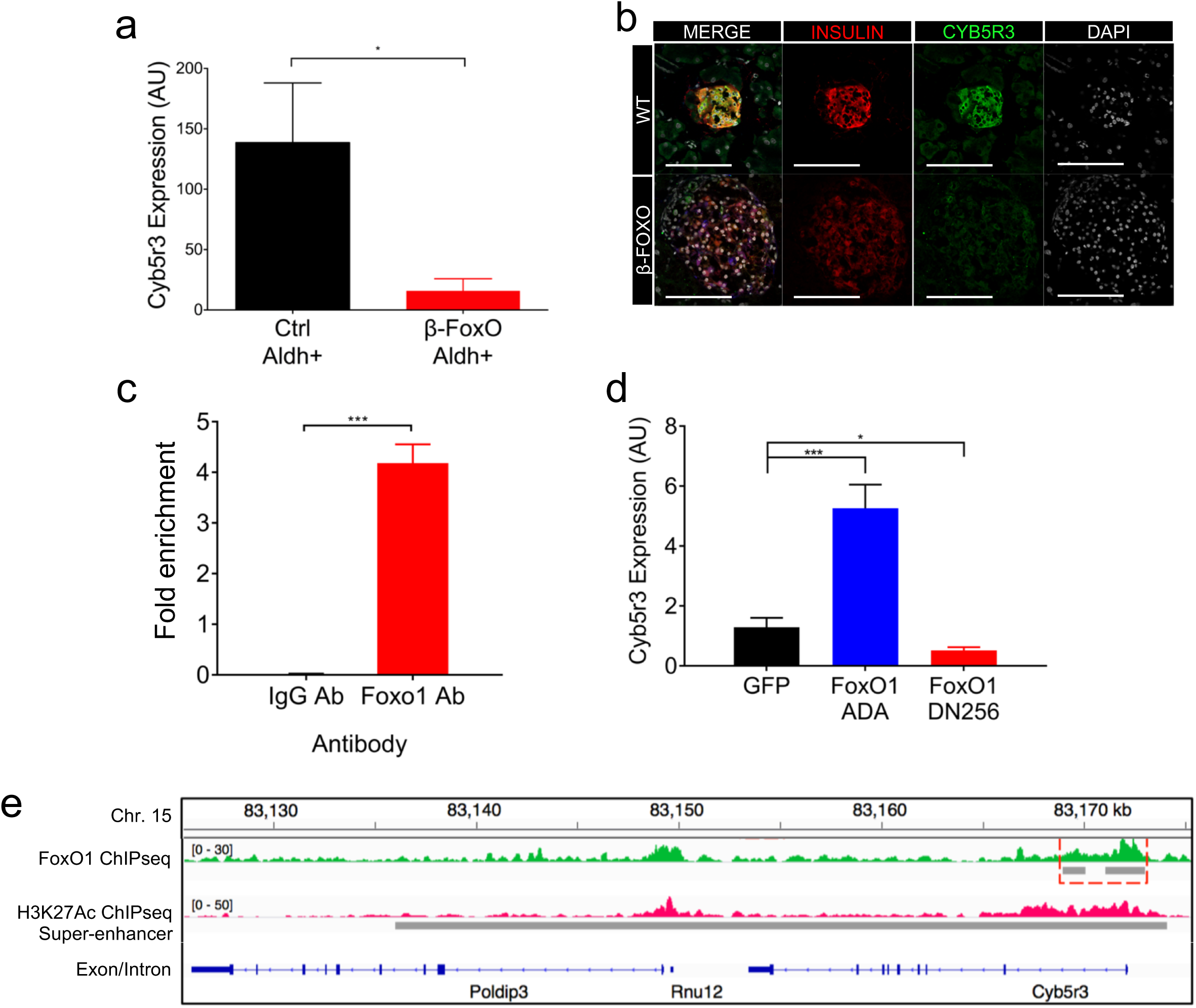
Regulation of β-cell Cyb5r3 by FoxO1. (A) *Cyb5r3* mRNA levels in β-cells with elevated Aldh activity (Aldh^hi^) from RIP-Cre^+^ (Ctrl) vs. RIP-Cre^+^ FoxO1/3/4^fl/fl^ (β-FoxO) mice (n=5 per group). (B) Immunofluorescence of Insulin (red), Aldh1a3 (blue), Cyb5r3 (green), and DAPI (white) in control vs. β-FoxO mice. (C) *Cyb5r3* promoter ChIP-qPCR with anti-FoxO1 or IgG control in Min6 cells. (D) *Cyb5r3* expression in Min6 cells transduced with adenovirus expressing GFP, constitutively active FoxO1-ADA, or dominant negative FoxO1-DN256. (E) Tracks of chromosome 15 FoxO1 (green) and H3K27ac ChIP-Seq (fuchsia) proximal to *Cyb5r3*. FoxO1 and super-enhancer sites are indicated by the gray bars. All data are presented as means ± SEM. *p < 0.05, **p < 0.01, ***p < 0.001 by Student’s t-test. All experiments were performed at least three times unless otherwise indicated.

We tested whether *Cyb5r3* is a FoxO1 target. Chromatin immunoprecipitation (ChIP) in mouse insulinoma (Min6) cells with an anti-FoxO1 antibody showed enrichment at a putative FoxO1 binding site (5’-ATAAACA-3’, –661 to –667) in the *Cyb5r3* promoter (Fig. 1C). To assess the effect of FoxO1 on *Cyb5r3* expression in β-cells, we transduced Min6 cells with adenovirus encoding constitutively active (FoxO1-ADA) or dominant negative (FoxO1-DN256) FoxO1 [32]. The former increased *Cyb5r3* expression ∼5-fold, while the latter suppressed it by 60% (Fig. 1D). We analyzed the endogenous *Cyb5r3* gene on chromosome 15 by ChIPseq of flow-sorted β-cells following immunoprecipitation with either anti-FoxO1 or anti-histone H3K27Ac to map regions of active chromatin [26]. We detected a strong enrichment of FoxO1 binding to the *Cyb5r3* promoter (Fig. 1E, green track). Furthermore, H3K27Ac ChIPseq revealed a super-enhancer associated with *Cyb5r3*, suggesting a regulatory role of this locus in β-cells (Fig. 1F, purple track) [33]. These data indicate that FoxO1 can directly regulate *Cyb5r3* expression.

### Cyb5r3 knock-down affects **β**-cell mitochondrial and secretory function

To determine whether Cyb5r3 is required for β-cell function, we transduced Min6 cells with adenovirus encoding a short hairpin RNA against Cyb5r3 (Ad-shCyb5r3). The shRNA lowered *Cyb5r3* mRNA and protein by 95% and 80%, respectively, while decreasing the expression of the related isoform Cyb5r4 by ∼30% (Supplementary Fig. 1A-D). When we assessed glucose-stimulated insulin secretion, cells transduced with shCyb5r3 adenovirus showed impaired insulin secretion compared to cells transduced with control adenovirus (Fig. 2A). Because Cyb5r3 is thought to participate in mitochondrial function [19], we measured basal respiration and observed a ∼25% decrease in Min6 cells stably expressing shCyb5r3 (Fig. 2B). Knockdown of the related isoform Cyb5r4 decreased mitochondrial respiration to a greater extent (∼40%).

**Figure 2.**
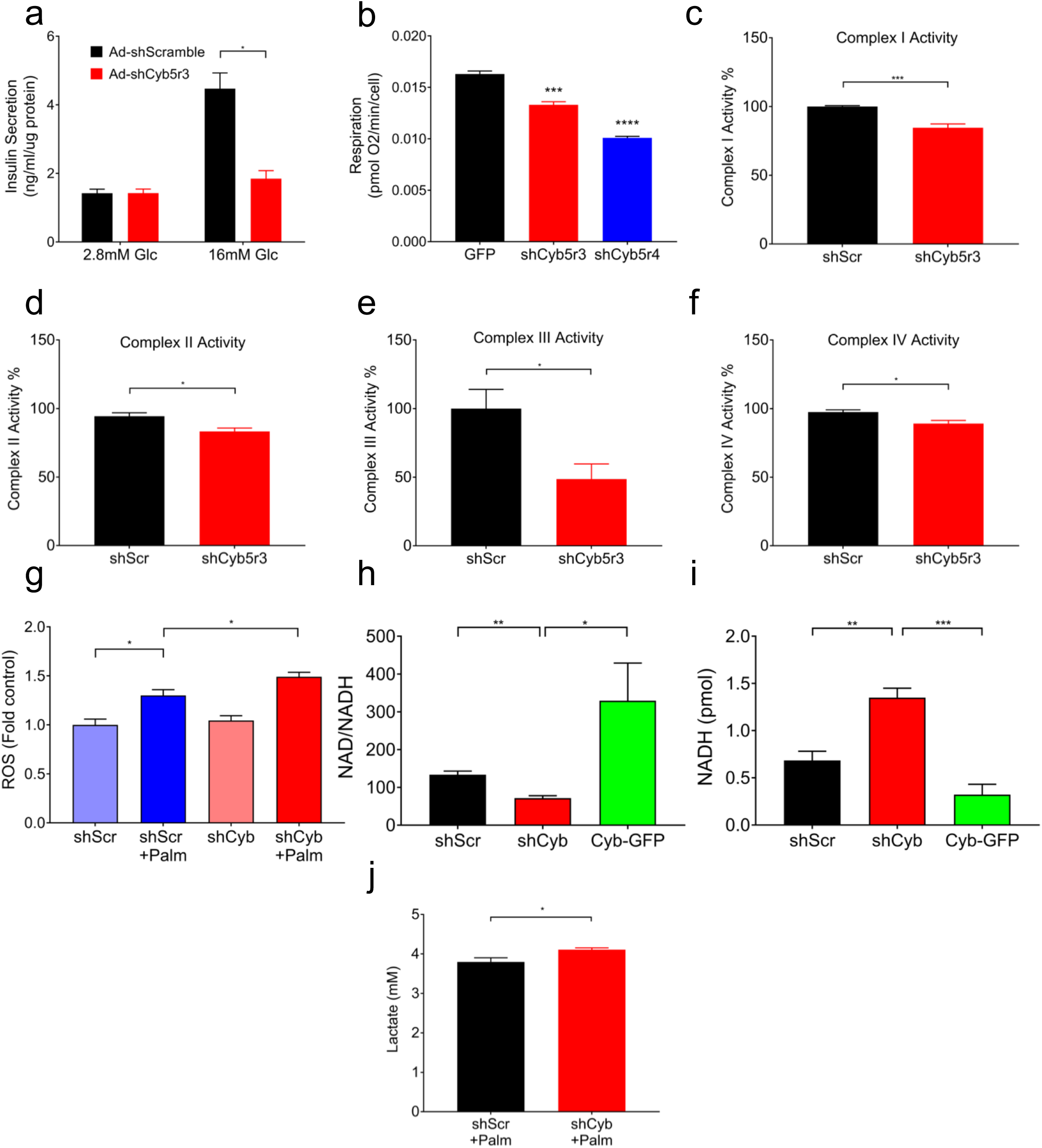
Cyb5r3 Regulates β-cell Secretory and Mitochondrial Function. (A) Glucose-induced insulin secretion in Min6 cells transduced with Ad-shCyb5r3 or Ad-shScramble. (B) Basal respiration in Min6 cells stably expressing GFP, shCyb5r3, or shCyb5r4. (C-F) ETC complex I-IV activity in mitochondrial fractions, (G) ROS levels in the presence or absence of 0.5mM palmitate, (H) NAD/NADH ratios, (I) NADH levels, and (J) Lactate levels in Min6 cells transduced with Ad-shCyb or Ad-shScr. All data are presented as means ± SEM. *p < 0.05, **p < 0.01, ***p < 0.001 by Student’s t-test. All experiments were performed at least three times unless otherwise indicated.

The second phase of insulin secretion in response to glucose is linked to mitochondrial generation of second messengers [34]. Although Cyb5r3 can affect mitochondrial ETC activity, the mechanism by which it does so is unclear [19; 21]. It can alter NADH availability for electron transfer, pass reducing equivalents to coenzyme Q, or reduce cytochrome b subunits or heme groups of ETC complex III. Thus, we sought to determine whether loss of Cyb5r3 activity in β-cells affects ETC activity through complex III, or whether it had a broader impact on all ETC complexes. Enzymatic assays for complexes I-IV revealed that mitochondria isolated from Ad-shCyb5r3-transduced Min6 cells had 4-15% reduction of complex I, II, and IV activity, and a much more severe ∼50% decrease of complex III activity (Fig. 2C-F), suggestive of a primary effect of Cyb5r3 on the latter.

We have proposed that a constitutive increase in β-cell lipid oxidation paves the way for β-cell failure and dedifferentiation, possibly by increasing ROS formation [17; 18]. ETC complexes I and III are the major sources of mitochondrial ROS production in states of stymied electron flow [35]. Because complex III (and to a lesser extent complex I) activity was decreased by shCyb5r3, we tested whether the latter also influenced lipid-induced ROS formation. To do this, we measured ROS in Ad-shCyb5r3-transduced Min6 cells cultured with palmitate, a condition that increases fatty acid oxidation and mimics the metabolic inflexibility of failing β-cells [17]. Consistent with our hypothesis, palmitate-treated Ad-shCyb5r3 cells had greater ROS production compared to controls (Fig. 2G).

As an NADH-dependent oxidoreductase, Cyb5r3 loss of function can perturb cellular NAD: NADH ratios. Min6 cells transduced with Ad-shCyb5r3 showed decreased NAD: NADH ratio, while overexpression of Cyb5r3-GFP led to an increase, consistent with Cyb5r3’s NADH-consuming activity (Fig. 2H, I). Furthermore, we hypothesized that upon knockdown of Cyb5r3, β-cells would compensate for increased NADH concentrations (or decreased NAD: NADH ratio) by increasing lactate production via lactate dehydrogenase to convert excess NADH to NAD^+^. This would be consistent with the increase in non-oxidative glycolysis and lactate dehydrogenase seen in diabetic islets [36]. Indeed, Ad-shCyb5r3-transduced Min6 cells showed increased lactate levels when cultured in 0.5mM palmitate (Fig. 2J). We suggest that exposure to palmitate stimulates fatty acid oxidation and generation of NADH, further necessitating lactate dehydrogenase-mediated conversion of NADH to NAD^+^. In the setting of increased lipid oxidation, the disturbances in NAD: NADH ratio, combined with increased ROS production and reduced electron transport, can induce cellular stress and β-cell failure.

### Cyb5r3 knock-down impairs insulin secretion and calcium influx *ex vivo*

Next, we tested the function of Cyb5r3 in primary islets by transducing them with shCyb5r3 or shScramble adenoviruses, followed by glucose-stimulated insulin secretion assays. Consistent with findings in Min6 cells, islets transduced with Ad-shCyb5r3 showed blunted insulin secretion in response to glucose despite normal insulin content. Interestingly, there was a trend toward higher basal secretion (Fig. 3A, B, and Supp. Fig. 1E).

**Figure 3.**
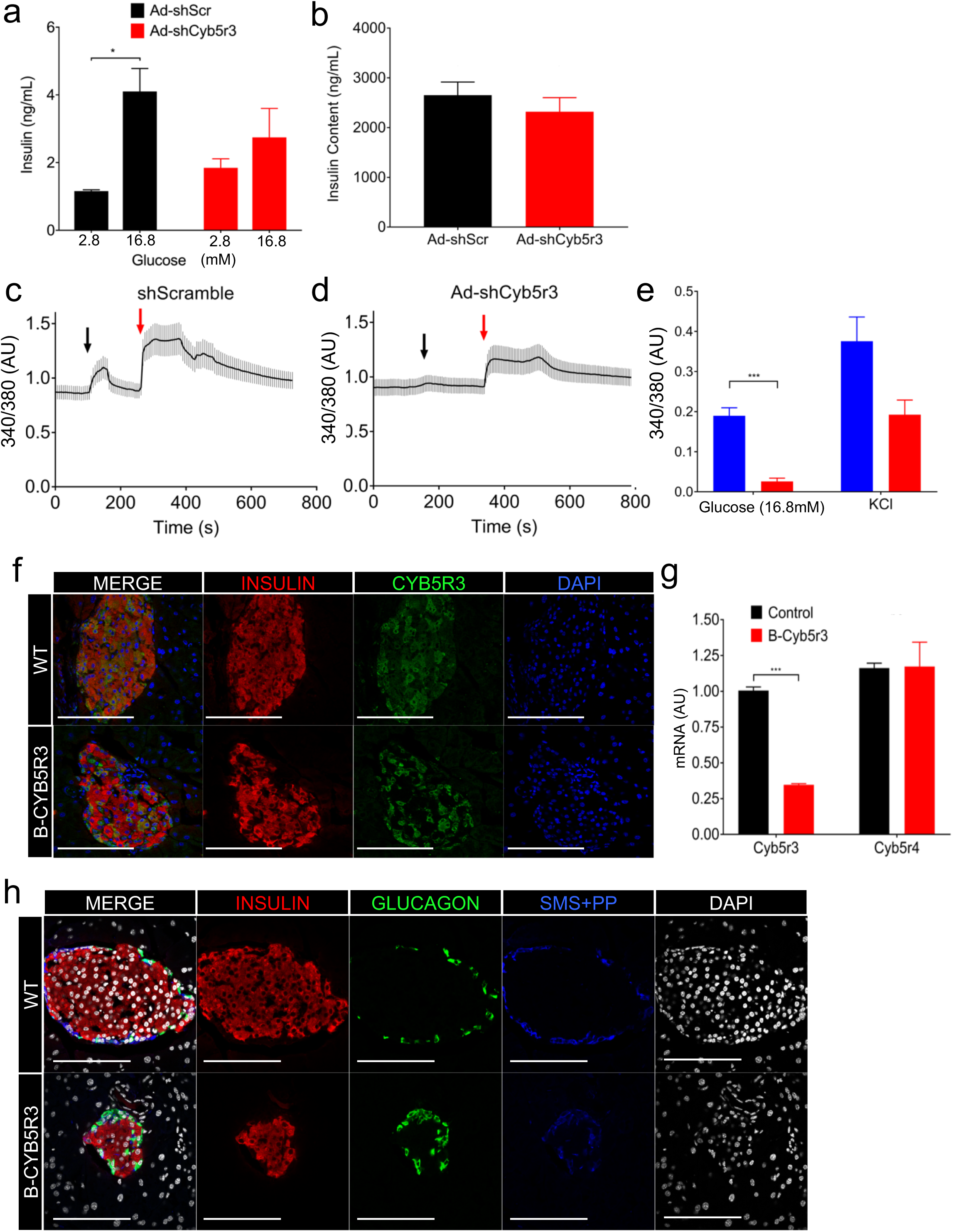
Cyb5r3 Regulation of Insulin Secretion *Ex vivo*. (A) Glucose-stimulated insulin secretion and (B) insulin content in wildtype islets transduced with Ad-shCyb5r3 or Ad-shScr (n=3 per group). (C) Representative calcium level traces measured by Fura2AM fluorescence in Insulin2-Gfp primary β-cells transduced with Ad-shCyb5r3 or Ad-shScr (n=10-20 per group). Black arrow denotes point at which perfusion changed from 2.8mM to 16.8mM glucose. Red arrow denotes addition of 40mM KCl. (D) Averaged maximal peak heights of the 340/380 ratio from the same experiment as in (C) (n=3 plates per treatment, with 10-30 beta cells counted per plate). (E) Insulin (red), Cyb5r3 (green), and DAPI (blue), (F) Insulin (red), Glucagon (green), Somatostatin with Pancreatic Polypeptide (blue), and DAPI (white) immunostaining in 2-month-old B-Cyb5r3 vs. RIP-Cre^+^ control mice (n=3 per group). All data are presented as means ± SEM. *p < 0.05, **p < 0.01, ***p < 0.001 by Student’s t-test.

To examine whether loss of Cyb5r3 impinges on intracellular calcium release, we measured Fura2AM calcium flux in single, chemically identified β-cells isolated from *Insulin2*-*GFP* mice [37] exposed to glucose or KCl. Following transduction with Ad-shCyb5r3 or Ad-shScramble, we gated single β-cells by GFP-expression for fluorescence measurements. Remarkably, Ad-shCyb5r3-transduced β-cells displayed a severely blunted calcium flux in response to glucose and, to a lesser extent, KCl (Fig. 3C-E).

### Impaired Insulin Secretion in **β**-cell-specific Cyb5r3 Knockout Mice

Next, we generated RIP-Cre-Cyb5r3*^fl/fl^*mice (B-Cyb5r3). These mice were born at Mendelian ratios without morphological defects or differences in body weight or composition (Supplementary Fig. 2A). Immunostaining of pancreata from female B-Cyb5r3 mice revealed a decreased number of Cyb5r3-immunoreactive β-cells (Fig. 3F) that dovetailed with a ∼65% decrease in *Cyb5r3* expression (Fig 3G). Notably, Cre recombination was much less efficient in male mice, and some islets retained Cyb5r3 expression in a majority of β-cells (Supplementary Fig. 3A). We do not know the reasons for this difference. Islets from 2-month-old B-Cyb5r3 mice had normal β and δ/pp cell content, but a greater proportion of α-cells compared to controls (17% vs. 11%, p<0.05) (Fig. 3H, Supplementary Fig. 4A).

We analyzed the metabolic features of B-Cyb5r3 mice. 4-month-old females demonstrated glucose intolerance relative to RIP-Cre^+^ controls as assessed by intraperitoneal glucose tolerance tests (IPGTT), as well as lower refed insulin levels after a 4-hour fast (Fig. 4A,C). Male mice showed a similar trend that did not reach statistical significance, possibly as a result of partial recombination (Fig. 4B). However, by 8 months of age, male mice also showed glucose intolerance compared to controls (Supplementary Fig. 3B). We observed similar results in oral glucose tolerance tests (OGTT), effectively ruling out an incretin effect (Supplementary Fig. 2C,D). In contrast, insulin tolerance tests (Fig. 4D,E), fasting or re-fed free fatty acids, and triglyceride levels were normal (Supplementary Fig. 2E,F).

**Figure 4.**
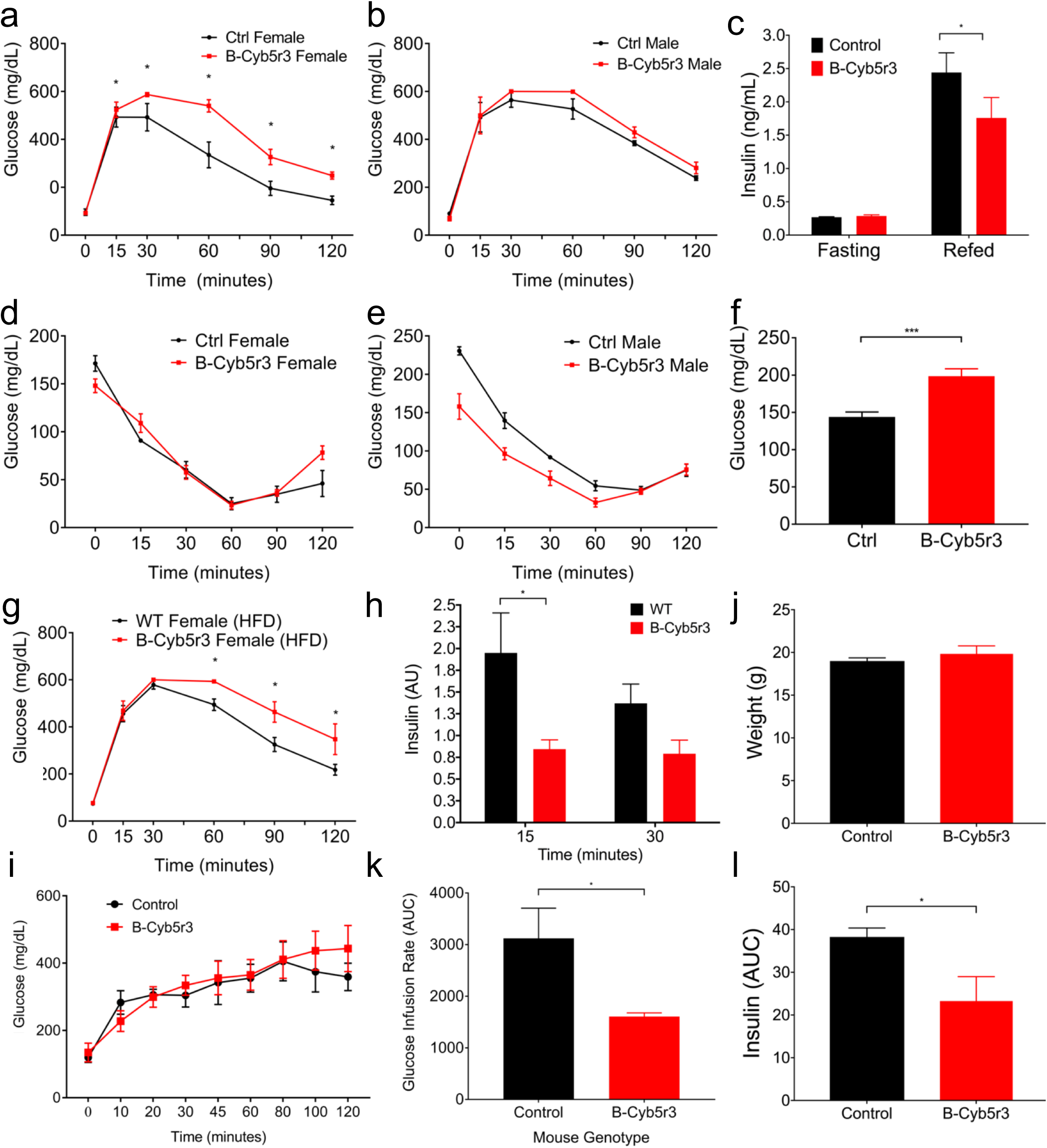
In vivo phenotyping of B-Cyb5r3 Mice. (A) Intraperitoneal glucose tolerance test (IPGTT) in 4-month-old female and (B) male B-Cyb5r3 vs. RIP-Cre^+^ mice. (C) 4-hr fasting and 1-hr refed insulin levels in female mice. (D) Insulin tolerance test in 4-month-old female and (E) male mice. (F) Glucose levels in female mice fed HFD for 1 week. (G) IPGTT in 4-month-old female mice fed HFD for 8 weeks. (H) Insulin levels at 15 and 30 minutes during IPGTT normalized by fasting levels in 4-month-old B-Cyb5r3 vs. RIP-Cre^+^ mice fed HFD for 8 weeks. (I) Glucose, (J) weight, (K) AUC of glucose infusion rate, and (L) AUC of serum insulin during hyperglycemic clamps in female B-Cyb5r3 vs. RIP-Cre^+^ mice. All data are presented as means ± SEM. *p < 0.05, **p < 0.01, ***p < 0.001 by Student’s t-test. (n=4 per group in experiments in A-D, and n=5 per group in experiments in F-L).

Next, we tested the effects of diet on B-Cyb5r3 female animals. (We did not study male animals in this experiment due to incomplete recombination.) 2-month-old B-Cyb5r3 mice placed on a diet consisting of 60% fat (HFD) developed hyperglycemia within a week (Fig. 4F). After 8 weeks of HFD, they showed glucose intolerance (Fig. 4G) and impaired insulin secretion after OGTT (Fig. 4H).

To assess insulin secretory function *in vivo*, we performed hyperglycemic clamps. Weight-matched female B-Cyb5r3 and wildtype controls were clamped at similar glucose levels (Fig. 4I,J). B-Cyb5r3 mice required a ∼50% lower glucose infusion rate (Fig. 4K) to maintain hyperglycemia as a result of a commensurate ∼40% decrease in plasma insulin levels (Fig. 4L), consistent with a primary insulin secretory defect.

### Functional and Ultrastructural Defects in Cyb5r3-deficient **β**-cells

We analyzed islet composition and markers of β-cell functional maturity. 2-month-old B-Cyb5r3 mice showed decreased levels of Glut2 and MafA (Fig. 5A,B), but preserved expression of Pdx1 (Fig. 5C). However, in 6-month-old B-Cyb5r3 animals, islet architecture was disrupted with α− and δ/pp-cells appearing in the islet core (Fig. 6A), and a greater relative proportion of α-cells (Supplementary Fig. 4B). In addition to Glut2 and MafA (Fig. 6B,C), the majority of β-cells now showed decreased Pdx1 expression (Fig. 6D). In addition, consistent with the incipient hyperglycemia, there was increased nuclear FoxO1 staining (Fig. 6E). It should be noted that this nuclear FoxO1 failed to restore expression of MafA, as predicted from prior work [14]. This suggests that the β-cell’s redox status or mitochondrial function participates in this process. We suggest that Cyb5r3-deficient β-cells attempt to compensate for impaired function by increasing FoxO1 activity, but since they lack Cyb5r3, they are unable to do so.

**Figure 5.**
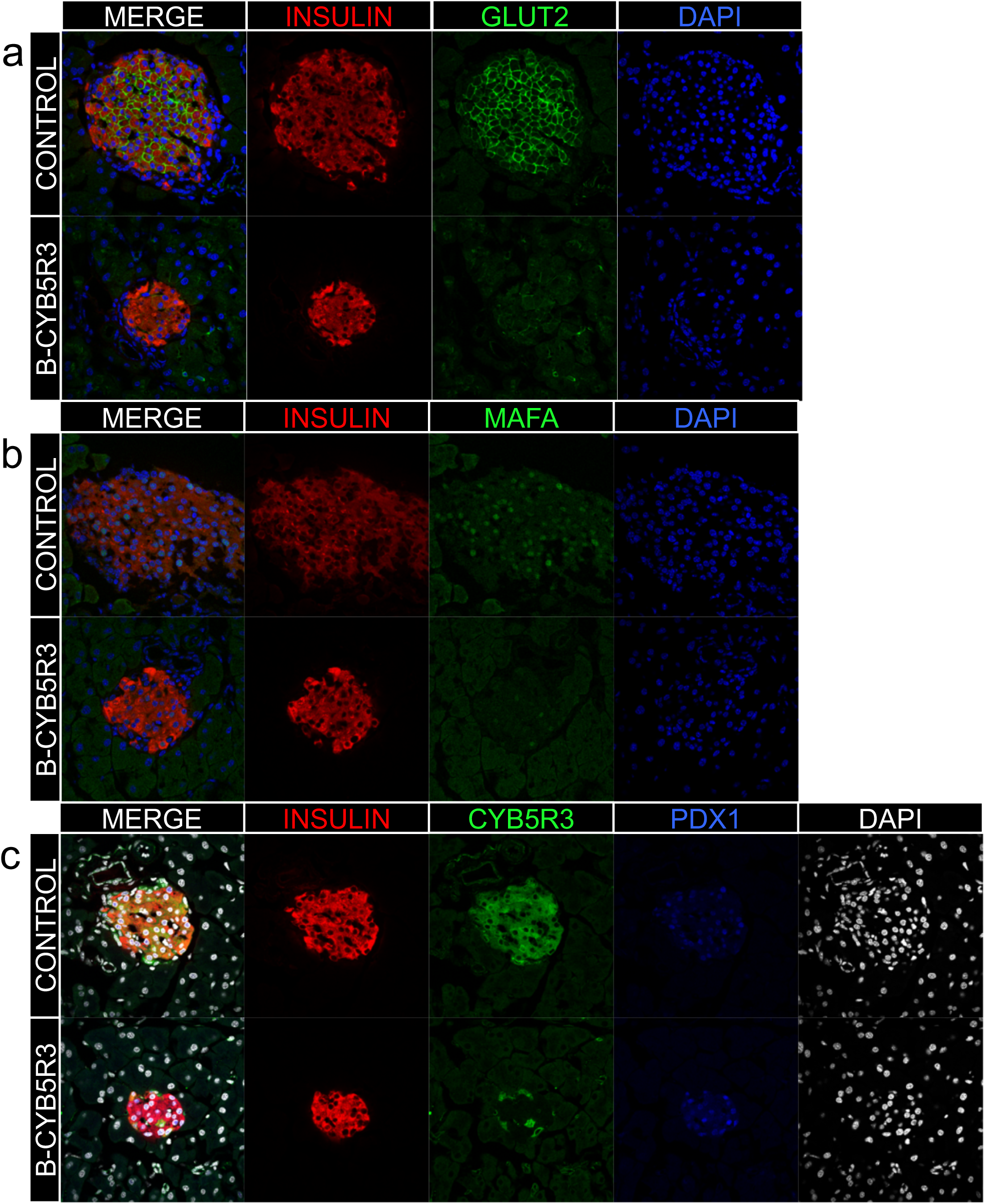
β-cell Markers in 2-month-old B-Cyb5r3 Mice. Immunostaining of pancreas sections from 2-month-old female B-Cyb5r3 vs. RIP-Cre^+^ control mice. (A) Insulin (red), Glut2 (green), and DAPI (blue). (B) Insulin (red), MafA (green), and DAPI (blue). (C) Insulin (red), Cyb5r3 (green), Pdx1 (blue), and DAPI (white) (n= 3 per group).

**Figure 6.**
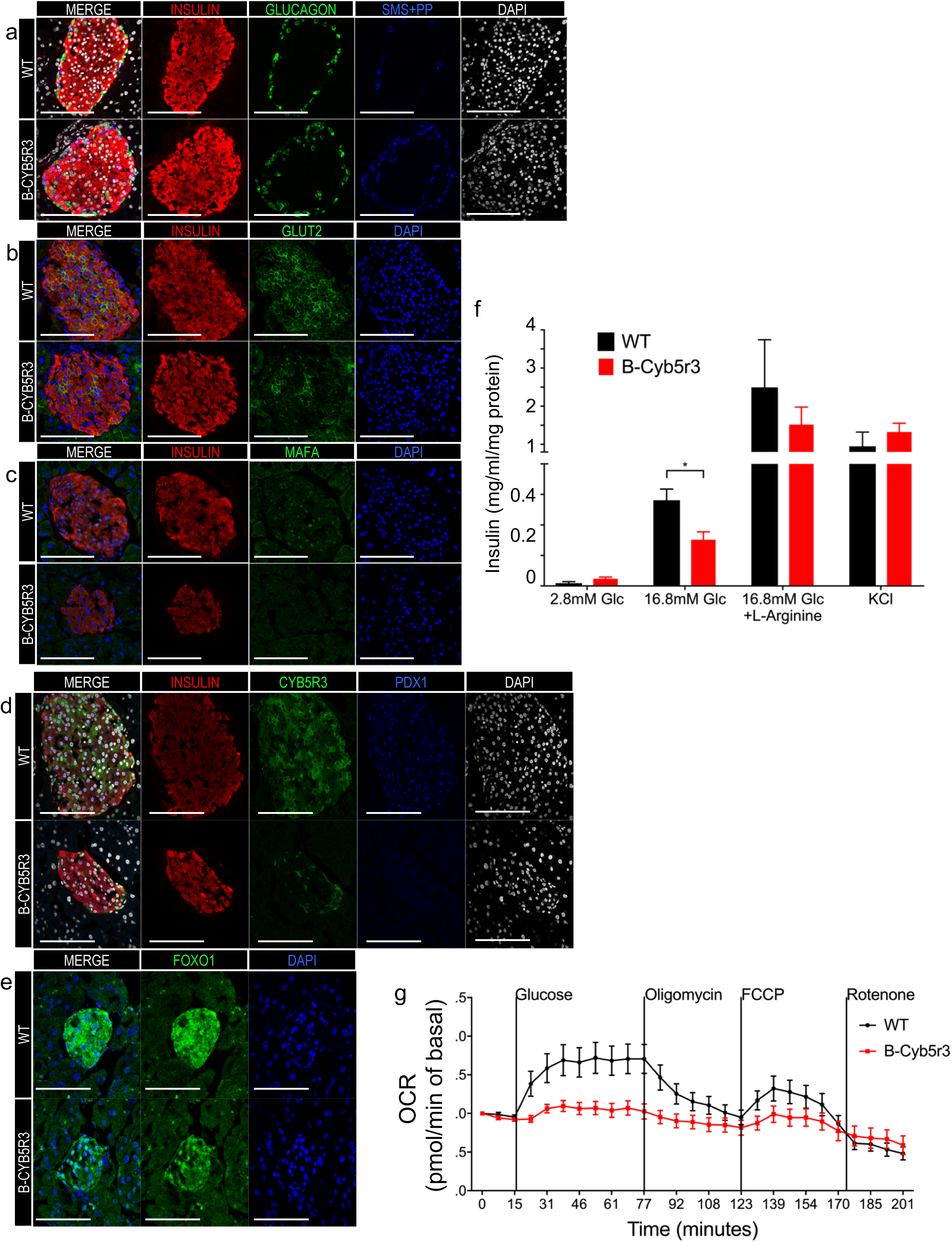
β-cell Markers and Insulin Secretion in 6-month-old B-Cyb5r3 Mice. Immunostaining of pancreas sections from 6-month-old female B-Cyb5r3 vs. RIP-Cre^+^ control mice. (A) Insulin (red), Glucagon (green), Somatostatin with Pancreatic Polypeptide (SMS+PP) (blue), and DAPI (white). (B) Insulin (red), Glut2 (green), and DAPI (blue). (C) Insulin (red), MafA (green), and DAPI (blue). (D) Insulin (red), Cyb5r3 (green), Pdx1 (blue), and DAPI (white). (E) FoxO1 (green) and DAPI (blue) (n=3 per group). (F) Glucose- and (G) secretagogue-stimulated insulin secretion in B-Cyb5r3 vs. RIP-Cre^+^ islets (n=4 samples each containing 5 islets from 3 mice for GSIS). (G) Mitochondrial respiration in B-Cyb5r3 vs. RIP-Cre^+^ islets. Vertical lines indicate timing of addition of glucose (25mM), Oligomycin (ATP synthase inhibitor), FCCP (uncoupler), and rotenone (complex I inhibitor). Islets from 5 mice were pooled in n=9 wells per genotype, with each well containing ∼70 islets. All data are presented as means ± SEM. *p < 0.05, **p < 0.01, ***p < 0.001 by Student’s t-test.

We also isolated primary islets from B-Cyb5r3 mice and performed glucose-stimulated insulin secretion assays. We observed decreased insulin secretion in response to glucose compared to RIP-Cre^+^ controls, but a normal response to L-arginine and KCl (Fig. 6F). To determine whether this impairment occurred in the context of mitochondrial dysfunction, we measured islets respiration (Fig. 6G). In WT islets, raising glucose levels from 5 to 25mM increased O_2_ consumption by ∼70%. In contrast, B-Cyb5r3 islets failed to respond, potentially linking decreased insulin secretion with impaired substrate-driven ATP production in B-Cyb5r3 mice. In summary, the data show that lack of Cyb5r3 results in multiple β-cell abnormalities and impairs insulin secretion.

To corroborate the above results, we analyzed β-cell ultrastructure by transmission electron microscopy. Control β-cells showed abundant mature insulin-containing granules with an electron-dense core (Fig. 7A), mitochondria with well-defined cristae (Fig. 7B,C, arrowheads), and well-organized endoplasmic reticulum (Fig. 7D, arrowheads). In contrast, B-Cyb5r3 β-cells had fewer mature insulin granules, accumulation of pale clear electron-dense vesicles lacking the typical sub-membrane clear halo and compatible with immature granules (Fig. 7E), and altered mitochondria with reduced cristae (Fig. 7F,G, arrowheads), often containing only traces of mitochondrial matrix precipitates (Fig. 7H, arrow). Additional alterations included endoplasmic reticulum stress as evidenced by the presence of numerous swollen cisternae (Fig. 7H, arrowheads) and incompletely formed secretory vesicles, suggestive of defects in granule assembly and maturation.

**Figure 7.**
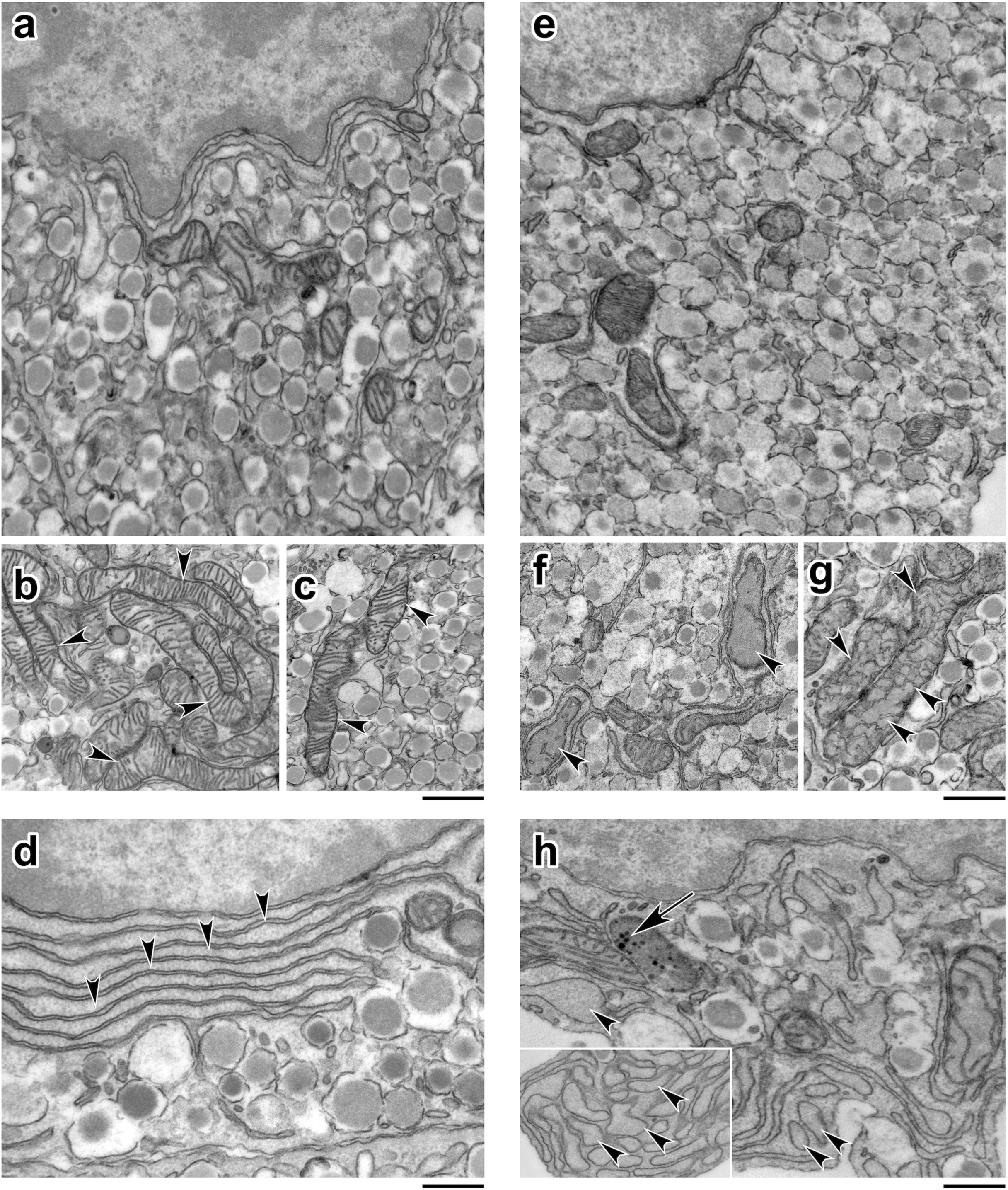
Transmission electron microscopy. (A-D) WT β-cells with secretory vesicles containing electron-dense core granules (A), mitochondria with typical cristae (B, C, arrowheads), and endoplasmic reticulum (D, arrowheads). (E-H) B-Cyb5r3 knockout β-cells with increased clear electron-dense immature vesicles (E), abnormal mitochondrial cristae (F-H, arrowheads), with occasional proteinaceous matrix precipitates (H, arrow), and swollen cisternae (H, arrowheads). Reference bar in A, B, C, E, F, G = 250 nm; in D and H = 100 nm. (n=3 mice and 50 cells per group).

## Discussion

Of the several pathways of β-cell failure [2], this work focuses on the FoxO1-dependent one. We have proposed the following model to explain β-cell failure in type 2 diabetes: in response to an altered metabolic milieu–brought about by excessive/inappropriate nutrition, genetic susceptibility, smoldering inflammation, or hormonal imbalance–β-cells activate a response network orchestrated by, but not limited to, FoxO1, whose mechanistic goal is to preserve β-cells from excessive mitochondrial lipid oxidation [38]. This compensatory mechanism fails if the inciting factor is not removed, because the FoxO1 response is self-limiting through stress-induced deacetylation-dependent degradation of the protein [14]. The loss of FoxO1, observed in common rodent models of diabetes such as *db/db* and HFD [7; 27; 29; 30], impairs mitochondrial oxidative phosphorylation, ATP production, and insulin secretion. This leads to dedifferentiation as a mechanism to prevent outright cellular death, because replacing β-cells by replication or neogenesis is at best an ineffective process [1].

However, the mechanistic link between FoxO1, mitochondrial failure, and β-cell dysfunction remained unknown. Although FoxO1 is known to regulate mitochondrial genes in response to insulin signaling in liver [39], the mechanism differs in β-cells, in view of the key role of glucose in driving mitochondrial function, and also because β-cell mitochondria are more uncoupled than liver’s [40]. The work described herein fills this gap in knowledge by identifying a FoxO1-dependent pathway through Cyb5r3 that regulates mitochondrial ATP production and prevents generation of toxic byproducts of mitochondrial oxidative phosphorylation, such as ROS, in response to fatty acids. Key features set Cyb5r3 apart from other mitochondrial oxidoreductases, including the closely related Cyb5r4 [22]: (*i*) Cyb5r3 expression parallels FoxO1’s, consistent with an effector function downstream of an important β-cell maintenance pathway; (*ii*) Cyb5r3 levels decrease as a function of insulitis progression in type 1 diabetic NOD mice [41] and in response to glucose toxicity in Ins1 cells [42]; (*iii*) Cyb5r3 regulates mitochondrial complex III, a critical site of ROS formation in response to lipids [40]; (*iv*) in this regard, it should be noted that failing β-cells (as defined by metabolic inflexibility and expression of dedifferentiation markers) present with selective dysfunction of complexes I, IV, and V [18]. Thus, additional complex III dysfunction in this context may represent the proverbial straw that breaks the β-cell’s back. Unlike complexes I/II, complex III generates superoxide in the matrix and on the outer side of the mitochondrial inner membrane [40]. This is consistent with Cyb5r3’s location to the outer mitochondrial membrane, and suggests that lack of Cyb5r3 deprives mitochondria of antioxidants to neutralize ROS outside the matrix, precipitating mitochondrial failure. We propose that Cyb5r3 links loss of FoxO1 with an impairment of oxidative phosphorylation that sets the stage for dedifferentiation. Although we didn’t investigate the latter point in detail, it’s supported by the observation that, in the absence of Cyb5r3, FoxO1 is unable to maintain expression of MafA.

Of the five genes encoding CYB5R, R3 is the main isoform in human β-cells [23]. We have recently mapped a conserved islet super-enhancer in humans and mice to *CYB5R3* [26]. Of note, a SNP in the *CYB5R3* promoter is associated with fasting glucose levels (Suppl. Fig. 5) [25]. No changes in *CYB5R3* levels have been reported in human diabetic islets [43–45]. However, as pointed out by the IMIDIA collaboration, there is no consensus on a diabetic gene expression signature, in view of the large variations observed in different datasets [44]. In this regard, it should be noted that the vast majority of genes whose biological function in human islets is beyond question don’t show changes to mRNA levels in these databases (e.g., all MODY genes except PDX1, glucokinase, sulfonylurea receptor, insulin processing enzymes, and others) [43–45]. In addition, CYB5R3 is expressed in both α− and β-cells; and a key point of our theory is that CYB5R3 levels drop only in those β-cells with advanced cellular pathology. Thus, detecting changes in expression levels of CYB5R3 in islets is technically difficult. We plan to study the function of human CYB5R3 in future experiments using iPS-derived β-cells.

Mitochondria allow β-cells to couple nutrient metabolism with insulin secretion, and mitochondrial dysfunction is present in rodent models and in human patients with diabetes [46–50]. β-cells are susceptible to ROS damage because of their low levels of free-radical quenching enzymes superoxide dismutase, catalase, and glutathione peroxidase [51]. The mitochondrial genome, inner membrane proteins and lipids are especially vulnerable to oxidative damage due to their proximity to free radicals, and damage to these structures impairs ETC function [40].

The observed decrease in mitochondrial complex activity following deletion of β-cell Cyb5r3 can be due to a direct effect on the production of reducing equivalents, or an indirect effect via the assembly of the ETC, as Cyb5r3’s flavin domain is known to reduce heme groups which are found in the cytochrome b and cytochrome c subunits of complex III, as well as in complex IV [52]. In addition, Cyb5r3 has been implicated in redox reactions [19]. Its yeast orthologue, NADH-Coenzyme Q reductase 1 (NQR1), reduces fasting-induced apoptosis by increasing levels of reduced Coenzyme Q and extends lifespan [53]. Yeast NQR1 activity increases in calorie restriction, a condition usually accompanied by increased fatty acid oxidation. Mice with Cyb5r3 gain-of-function have increased complex I and III, and unsaturated very long chain FAs, as well as reduced ROS [21]. In mammals, Cyb5r3 also localizes to the endoplasmic reticulum, where it has been proposed to catalyze fatty acid desaturation/ elongation, cholesterol biosynthesis, and xenobiotics metabolism. Thus, the abnormalities of secretory vesicle formation and maturation in Cyb5r3-deficient β-cells can alternatively result from effects of Cyb5r3 on cholesterol biosynthesis [54]. In addition to FoxO1, Cyb5r3 expression can be stimulated by a complex of FoxO3a and Nrf2 in response to oxidative stress [19]. Abnormalities of Nrf2 function result in β-cell dysfunction [55] and impaired Nrf2 expression is found in FoxO-deficient β-cells [17].

In summary, this work identifies an effector of FoxO1 loss-of-function-associated β-cell failure, providing new insight into this process.

## Methods

### Animals

Mice were housed under standard conditions (lights on at 07:00 and off at 19:00) and fed normal chow (NC) (PicoLab rodent diet 20, 5053; Purina Mills). NC had 62.1% calories from carbohydrates, 24.6% from protein and 13.2% from fat. High fat diet (HFD) had 20% calories from carbohydrates, 20% from protein and 60% from fat (D12492; Research Diets). Both genders of 2- to 60-day-old mice were used for developmental studies. Female B-Cyb5r3 mice aged 8-24 weeks were used except where indicated that we used both genders, due to less efficient Cre recombination in males B-. Genotyping was performed as previously described [37; 56-58]. To generate RIP-Cre^+^ Cyb5r3-floxed mice, RIP-Cre^+^ mice [59] were crossed with Cyb5r3-floxed mice (Knockout Mouse Project, Cyb5r3^tm1a(KOMP)Wtsi^) and were maintained on a C57BL/6J background (Jackson Laboratories #000664. The floxed *Cyb5r3* allele was genotyped using the primers 5’-ACAGTCCAGCTTTGGCTTTACCC-3’ and 5’-ATAGGGCTAGAAAAGGAGCAGAGAGC-3’ yielding a 456bp product. We derived Cre^+^ controls from the same litters. Sample size calculations were based on the variance observed in prior experiments [17; 18].

### Cell Lines

We used glucose-responsive Min6 insulinoma cells [14] cultured in DMEM (Invitrogen 41965-039) containing 15% FBS (Sigma-Aldrich F2442), penicillin-streptomycin (ThermoFisher 15140122) and 0.05mM 2-mercaptoethanol (Gibco 31350010) at 37°C and 5% CO_2_. The cell line has not been authenticated, but tested negative for mycoplasma by PCR-based detection kit (Sigma-Aldrich D9307). Medium containing 0.5mM palmitate was prepared by first dissolving FFA-free BSA (Fisher Scientific BP9704100) in Min6 media to a final concentration of 0.6 M, then adding sodium palmitate (Sigma-Aldrich P9767). The solution was shaken overnight at 250rpm at 50**°**C, and filtered before use.

### Primary Islet Culture

We isolated islets by collagenase digestion from female mice as described [17]. Collagenase P (Sigma 11249002001) was diluted in Medium 199 (ThermoFisher 11150067) to a concentration of 1mg/ml and kept on ice. The animal was sacrificed in a CO_2_ chamber followed by cervical dislocation. An abdominal incision was made, the common bile duct was clamped with a hemostat near the liver and 3ml of cold Collagenase P solution was injected into the hepatopancreatic ampulla to inflate the pancreas. The excised pancreas was incubated in a vial at 37°C with shaking 16min. Thereafter, vials were shaken and filtered through a stainless-steel strainer. Medium 199 was added to a final volume of 50ml, and the mixture centrifuged at 1,100rpm for 2min at 4°C. The supernatant was removed and 10ml of 4°C histopaque was added to the pellet. The mixture was vortexed and 10ml of Medium 199 was layered on top of the histopaque followed by centrifugation at 2,700rpm for 20min at 4°C. Islets at the interface of the histopaque and Medium 199 were removed and placed in 15ml tubes. Medium 199 containing 10% FBS (Sigma-Aldrich F2442) was added to a final volume of 15ml followed by centrifugation at 1,100rpm for 2min at 4°C. This step was repeated twice more. The pellet was then resuspended in 15ml Medium 199 + 10% FBS, and islets were allowed to settle on ice for 5min. This wash was repeated twice, and islets were handpicked into RPMI 1640 media (ThermoFisher 11150-067) containing 15% FBS (Sigma-Aldrich F2442) for further analysis.

### Adenoviral Vectors

Dominant negative FoxO1 (DN256) and constitutively active FoxO1 (FoxO1-ADA) adenovirus have been described [60]. shCyb5r3 adenovirus was generated by cloning the effective shRNA sequence (5’-GATTGGAGACACCATTGAAT-3’) into the pEQU6-vector with LR Clonase II (ThermoFisher 11791100), then ligating to pAd-REP. The purified cosmid DNA (2 μg) was digested with Pac1 and transfected into 293 cells with Lipofectamine 2000 (ThermoFisher 11668019). Adenovirus plaques appeared 7 days after transfection. The virus was amplified to 10^12 vp/ml using a cesium chloride gradient. shScramble adenovirus from Welgen (V1040) was generated using the same procedures.

### RNA Measurements

We isolated RNA with RNeasy Mini-kit (QIAGEN) and reverse-transcribed 1ug of RNA using qScript cDNA SuperMix (Quanta). cDNAs were diluted 1:5 and qPCR was performed using GoTaq® qPCR Master Mix (Promega). Primers for mCyb5r3 were: (F: 5’-CAGGGCTTCGTGAATGAGGAG-3’, R: 5’-TCCACACATCAGTATCAGCGG-3’). All other PCR primer sequences have been published [7]. Gene expression levels were normalized to hypoxanthine-guanine phosphoribosyltransferase (HPRT) using the ΔΔCT method and are presented as relative transcript levels.

### Protein Analysis

Min6 cells were lysed in ice-cold buffer (20mM Tris-HCl, pH7.4, 150mM NaCl, 10% glycerol, 2% NP-40, 1mM EDTA, 20mM NaF, 30mM Na4P2O7, 0.2% SDS, 0.5% sodium deoxycholate) supplemented with Protease/Phosphatase Inhibitor Cocktail (1X, Cell Signaling) and centrifuged for 10min (14,000rpm). Protein concentration was assessed by Pierce BCA protein assay (Thermo scientific). For Western Blotting, 30-50ug of protein were fractionated on gels. Densitometric analysis was performed using ImageJ (NIH).

### Chromatin Immunoprecipitation (ChIP) Assay

ChIP was performed as described [61]. Min6 cells were cross-linked, lysed, and sonicated. Immunoprecipitations were done with 4μg of anti-Foxo1 (Abcam Ab39670) or IgG. Immunocomplexes were recovered with protein G-dynabeads (Life Technologies 10003D), washed, and reverse-crosslinked. The precipitated DNA was analyzed by quantitative PCR using primers for mCyb5r3 (F: 5’-CATCTAGTGGAATGGGTACGTG-3’, R: 5’-TAGTGCAGAACGGTCTTTGTAG-3’). Fold-enrichment was calculated using a modified ΔCT method and normalized to DNA immunoprecipitated with IgG control.

### Fluorescence-Activated Cell Sorting of **β**-cells

We isolated mouse islets by collagenase digestion [17] and sorted β-cells as described [18]. Briefly, we incubated cells with the fluorescent ALDH substrate BODIPY™-aminoacetaldehyde (Aldefluor) for 1hr prior to flow cytometry. Thereafter, cells were loaded to a BD Influx sorter and analyzed with a BD LSRII. We gated cells for RFP (red) and aldefluor (green) fluorescence, yielding three sub-populations: RFP^−^ALDH^−^ (non-β-cells), RFP^+^ALDH^−^ (β-cells), and RFP^+^ALDH^+^ (ALDH-positive β-cells).

### RNA Sequencing of RIP-Cre^+^ FoxO1^fl/fl^**β** cells

RNA was isolated with Nucleospin RNA kit (Macherey-Nagel) with DNase I treatment. Directional poly-A RNAseq libraries were prepared and sequenced as PE100 (100bp paired-end reads) on Illumina Hiseq 2500 in high output mode. For analysis, we used 60 to 73 million read-pairs, and the TopHat algorithm to align reads to mouse genome mm10. Alignments in BAM files were further analyzed using Cufflinks software, followed by Cuffcompare and Cuffdiff. For statistical analysis, a standard cutoff of *p* < 0.05 was applied [62].

### ChIP Sequencing and Super-Enhancer Analysis

We isolated islets from male FoxO1-GFP knock in mice [26] and cultured them overnight in RPMI 1640 (Gibco) supplemented with 15% FBS (Corning). Cells were fixed with 1% formaldehyde for 10 min at room temperature. The reaction was quenched with 0.125 M glycine, cells were pelleted, washed with PBS, and frozen at −80°C. ChIP and ChIP libraries were performed as described using anti-GFP (Abcam ab290) or anti-H3K27ac (Active Motif, 39133) antibodies [26]. The resulting DNA libraries were quantified with Bioanalyzer (Agilent), and sequenced on Illumina NextSeq 500 with 75-nt reads and single end. Reads were aligned to mouse genome mm10 using the Burrows-Wheeler Aligner (BWA) algorithm with default settings. MACS (1.4.2) algorithm was used for peak calling with a standard cutoff of 1×10^-7^. ROSE was used to identify enhancers [33] and super-enhancers [63]. MACS peaks identified by H3K27ac ChIPseq were used as “constituent enhancers” input in ROSE [63]. Default settings for stitching distance (12.5 kb) and transcription start site exclusion zone (0 bp− no promoter exclusion) were used.

### Insulin secretion assays

We placed islets in ice-cold KRBH buffer (119 mM NaCl, 2.5 mM CaCl_2_, 1.19 mM KH_2_PO_4_, 1.19 mM Mg_2_SO_4_, 10 mM HEPES pH 7.4, 2% BSA and 2.8 mM glucose) and incubated at 37°C for 1hr (or overnight for Min6 cells), followed by the addition of varying glucose concentrations (2.8, 16.8 mM), or 30mM L-Arginine (Sigma-Aldrich A5006), or 40mM KCl (Sigma-Aldrich P9541) for 1hr at 37°C. At the end, we collected islets by centrifugation and assayed the supernatant for insulin by ELISA (Mercodia, #10-1247-01) [64; 65]. Insulin levels were normalized by protein or insulin content.

### Measurement of Mitochondrial Respiration

Mitochondrial respiration was measured using the XF24e Extracellular Flux Analyzer (Seahorse Bioscience). For cultured cells, oxygen consumption was sequentially measured under basal conditions (Seahorse media with 10mM glucose and 2mM pyruvate) and following addition of 1μM oligomycin (complex V inhibitor), 0.75μM carbonyl cyanide-p trifluoromethoxyphenylhydrazone (FCCP) and 1μM rotenone/1μM antimycin A (complex I and complex III inhibitors, respectively). For islets, we used XF24 islet capture microplates in Modified XF Assay Media (MA) (supplemented XF DMEM assay media with 3mM glucose and 1% FBS). Islets were washed and concentrated in this supplemented medium. 70 islets were seeded in 100μl in each well and accommodated into the depressed well. Following addition of the individual screens, wells were supplemented with 400μl MA media. Oxygen consumption was measured under basal conditions (MA) and after the sequential addition of 20mM glucose, 10μM oligomycin, 3μM FCCP and 5μM rotenone. Prior to experiments, seahorse plates were preincubated at 37°C without CO_2_ for 1hr to equilibrate temperature and adjust metabolism. Min6 results represent averages of ≥3 biological replicates, each consisting of 4-5 technical replicates with 60,000 cells/well. Islet experiments represent 10 technical replicates of each genotype and results are shown as individual experiments. All oxygen consumption (OCR) data were normalized by cell number or protein content.

### Mitochondrial Complex Activity Assays

Complex I-IV activity was measured as described in mitochondrial-enriched fractions, with the following modifications for Complex I, II, and IV [66]. For Complex I activity, we used NADH dehydrogenase assay buffer (1X PBS, 0.35% BSA, 200μM NADH, 240μM KCN, 60μM DCIP, 70μM decylubiquinone, 25μM Antimycin A) containing 200μM Rotenone or water; for Complex II activity, succinate dehydrogenase assay buffer (25mM K3PO4 pH 7.2, 0.35% BSA, 2mM EDTA, 5mM sodium succinate, 60μM DCIP, 25μM Antimycin A, 240μM KCN, 2μM Rotenone, and 150μM phenazine methosulfate). Complex I and II activity were measured as rate of decrease in absorbance at 600nm. Specific Complex I activity was calculated by subtracting rotenone-resistant activity from total activity. For Complex IV activity, we used cytochrome C oxidase assay buffer (20mM K3PO4 pH 7.2, 0.35% BSA, 1mM EDTA, 25μM Antimycin A, 100μg/mL reduced cytochrome C) containing either 10mM KCN or water. Complex IV activity was measured as rate of decrease in absorbance at 550nm. Specific Complex IV activity was calculated by subtracting the KCN resistant activity from the total activity. For each assay, absorbance was measured at the given wavelength every 10 secs for 20 min at 25°C using a SpectraMax Paradigm Multi-Mode Microplate Reader (Molecular Devices LLC., Sunnyvale, CA). Enzymatic activities were normalized to protein concentrations. Enzyme activity was calculated as described [66].

### Reactive Oxygen Species (ROS)

Intracellular ROS was measured in Min6 cells by conversion of the acetyl ester CM-H_2_DCFDA into a fluorescent product (excitation/emission 485nm/520nm). Min6 cells were grown in complete medium in 24-well plates and washed with at Hanks buffered salt solution (HBSS). Immediately before the assay, 50ug of CM-H_2_DCFDA was reconstituted at 10mM in 8.6ul DMSO. The HBSS was then replaced with 0.8ml HBSS containing 1uM CM-H_2_DCFDA and incubated at 37°C for 30 min. The plate was placed on ice and wells were washed with cold HBSS. Cells were scraped in 0.5ml of cold HBSS and 200ul of the cell suspension was loaded into a black uncoated 96-well plate for reading. The remainder of the sample was used to assay for total protein to normalize cellular content per well.

### NAD^+^/NADH and lactate measurement

NAD^+^ and NADH levels were quantified using the BioVision K337 kit according to the manufacturer’s instructions. In brief, MIN6 cells were grown in complete medium (25mM glucose), then lysed by freeze/thaw. To determine NADH levels, samples were heated at 60°C for 30 min to decompose NAD. To detect total NAD, samples were incubated with the NAD cycling enzyme mix to convert NAD^+^ to NADH. Both samples were then mixed with NADH developer and incubated at room temperature for 1hr before colorimetric reading at 450 nm. NAD levels were calculated by subtracting NADH from total NAD. Lactate was measured using the Abcam kit (Ab65331). In brief, Min6 cells grown in complete medium (25mM glucose) were washed, homogenized, and the supernatant collected. Ice-cold perchloric acid (4M) was added to a final concentration of 1M. Samples were incubated on ice for 5 min and spun at 13,000 x g for 2min at 4°C. Following neutralization with KOH, the supernatant was collected for colorimetric assay at 450nm.

### Single-Cell Intracellular Calcium Microfluimetry

Islets from B-Cyb5r3 and control mice were allowed to recover for 2hr in RPMI medium containing 15% FBS and penicillin-streptomycin (Islet media). Islets were dispersed to single cells using trypsin in a 37**°**C water bath, plated on fibronectin-precoated 35mm glass bottom dishes with 20mm microwells (Sigma, F1141) and transduced with shScramble or shCyb5r3 adenovirus (100 MOI) in islet media. After overnight incubation at 37**°**C, cells were washed with islet media and allowed to recover for two days. On the third day, each plate was loaded in the dark with 5uM fura2-AM (ThermoFisher F1221) in KRBH buffer. Cells were washed, transferred to a perifusion chamber placed in the light path of a Zeiss Axiovert fluorescence microscope (Zeiss, USA), and perifused with glucose (2.8 or 16.8mM), or KCl (40mM)-containing KRBH buffer. β**-**cells were excited by a Lambda DG-4 150 Watt xenon light source (Sutter, Novato, USA), using alternating wavelengths of 340 and 380 nm at 0.5□s intervals, and imaged at 510 nm. For each data set, regions of interest corresponding to the locations of 10-20 individual cells were selected and digital images were captured using an AxioCam camera controlled by Stallion SB.4.1.0 PC software (Intelligent Imaging Innovations, USA). Single-cell intracellular Ca^2+^ mobilization data consisted of excitation ratios (F340/F380) plotted against time (min).

### Metabolic Analyses

We performed intraperitoneal glucose tolerance tests (ipGTT) (2 g/kg) after overnight fast, and intraperitoneal insulin tolerance tests (0.75 units/kg) after a 5-hr fast [65; 67]. Glucose was measured from tail vein using OneTouch (One Touch Ultra, Bayer). We measured serum insulin by ELISA (Mercodia #10-1247-01). We performed hyperglycemic glucose clamps [17] and measured body composition as described [68]. We measured non-esterified fatty acids with HR Series NEFA-HR(2) (Wako Pure Chemicals) and triglycerides with Infinity #TR22421 (ThermoFisher).

### Immunohistochemistry

We performed immunohistochemistry as described [7]. Pancreata were placed in 4% paraformaldehyde (PFA) at 4°C for 4hr, washed 3X with ice-cold PBS, and placed in 30% sucrose overnight. Tissue was embedded in Tissue-Tek® optimal cutting compound (Sakura® Finetek), frozen on dry ice, and cut into frozen 6-um-sections. Sections were air-dried for 20min before immunostaining. Antibodies are listed in supplementary Table 1. Images were captured using a Zeiss LSM 710 confocal microscope using a 45X objective and analyzed using ZEN.

### Transmission electron microscopy

Ultrastructural analysis of pancreatic islets from WT and in B-Cyb5r3 mice was performed as described [55]. Islets were fixed in 0.1 M sodium cacodylate buffer, pH7.4, 2% paraformaldehyde, 2.5% glutaraldehyde (Electron Microscopy Sciences), and 3 μM CaCl_2_. Samples were post-fixed with osmium tetraoxide (1% w/v in H_2_O) and counterstained with uranyl acetate (2% w/v in H_2_O). After embedding in Durcupan resin (Sigma-Aldrich), ultrathin sections (70 nm) were prepared, mounted on 300 mesh gold grids and counterstained with uranyl acetate (1% w/v in H_2_O) and Sato lead (1% w/v in H_2_O). Ultrathin sections were imaged at 80 keV using a JEOL JEM-1230 microscope, equipped with an AMT XR80 CCD camera.

### Statistics

Statistical analyses were performed using Prism 6.0/7.0 software (Graph Pad). We calculated p-values for unpaired comparisons between two groups by two-tailed Student’s t-test, using the customary threshold of *p* <0.05 to declare significance. One-way ANOVA followed by Tukey’s multiple comparisons test (compare all pairs of columns) was used for comparisons between three or more groups. Two-way ANOVA followed by Bonferroni post-test was used to examine two different variables. All results are presented as means ± SEM. Sample sizes were estimated from expected effect size based on previous experiments and can be found in the figure legends. No randomization or blinding was used.

### Data Availability

The datasets are available from the corresponding author on reasonable request.

### Study Approval

All animal experiments were in accordance with NIH guidelines for Animal Care and Use, approved and overseen by Columbia University Institutional Animal Care and Use Committee (IACUC).

## Author Contributions

J.F. and W.D. designed and performed experiments, analyzed data, and wrote the manuscript. J.Y.K.-M. designed experiments. J.S. performed the ChIP assay. T.K. performed RNA sequencing, ChIP sequencing, and acetylation mapping. D.L. performed Seahorse respirometry. C.G. performed mitochondrial complex activity assays. T.K. and M.K. performed mouse experiments. E.O. oversaw research. V.C. performed studies of β-cell ultrastructure by Transmission Electron Microscopy and analyzed data. D.A. designed experiments, oversaw research and wrote the manuscript.

## Acknowledgements

This work was supported by NIH grants DK112518 to J.F., DK64819 to D.A., DK103711, Program Grant 4553677 from the WA State Life Sciences Discovery Fund, Innovation Pilot Award from UW Institute for Stem Cell and Regenerative Medicine, and Washington State funds to V.C. We are grateful to members of the Accili laboratory and to Enrique Garcia (Columbia University) for insightful data discussions. We thank Thomas Kolar, Ana Flete-Castro, Jun B. Feranil, Lumei Xu, and Q. Xu (Columbia University) for outstanding technical support (DK63608). We thank Travis Morgenstern and Dr. Henry Colecraft (Columbia University) for training and equipment for calcium flux imaging, Dr. Magali Mondin (U. Nice) for the immunostaining quantification macro for ImageJ. We acknowledge the expert assistance of Edward Parker at the Vision Science Center (EY001730), University of Washington (Seattle, WA) for assistance with Transmission Electron Microscopy.

## Supplemental Information

**Supplementary Figure 1. Adenovirus-mediated shCyb5r3 Knockdown, Related to** Figure 2. (A) *Cyb5r3* in Min6 cells transduced with shCyb5r3 *vs*. shScramble at varying MOIs. (B) *Cyb5r4* mRNA in the same samples as (A). (C) Densitometry of Cyb5r3 protein levels in samples transduced with shScr vs. varying MOIs of shCyb5r3 adenovirus. (D) Western blot used for densitometry shown in (C). Rat Ins1 cells were used as a negative control for the mouse-specific shCyb5r3 sequence. (E) *Cyb5r3* mRNA in WT islets transduced with Ad-shScramble vs. Ad-shCyb5r3. All data are presented as means ± SEM. *p < 0.05, **p < 0.01, ***p < 0.001 by Student’s t-test. All experiments were performed at least 3 times in triplicate.

**Supplementary Figure 2. Metabolic Phenotyping of B-Cyb5r3 mice, Related to** Figure 4. (A-B) Body composition measured by MRI of B-Cyb5r3 vs. RIP-Cre^+^ mice fed (A) normal chow and (B) HFD. (C-D) Oral glucose tolerance test (OGTT) of 4-month-old B-Cyb5r3 (C) female and (D) male mice fed normal chow. (E) Fasting and refed triglycerides and (F) non-esterified free fatty acids (NEFAs) in B-Cyb5r3 mice on normal chow. All data are presented as means ± SEM. *p < 0.05, **p < 0.01, ***p < 0.001 by Student’s t-test (n=4-6 per group).

**Supplementary Figure 3. Cyb5r3 Expression and IPGTT in Male B-Cyb5r3 Mice, Related to** Figures 3 and 4. (A) Immunostaining of Insulin (red), Cyb5r3 (green), Pdx1 (blue), and DAPI (white) in male B-Cyb5r3 pancreata (n=3 per group). (B) IPGTT in 8-month-old male B-Cyb5r3 mice vs. RIP-Cre^+^ controls (n=4 per group). All data are presented as means ± SEM. *p < 0.05, **p < 0.01, ***p < 0.001 by Student’s t-test.

**Supplementary Figure 4. Islet Cell Quantification, Related to** Figures 3 and 6. (A) Insulin, Glucagon, and Somatostatin + PP area normalized to total islet area as determined by immunohistochemistry. Data shown for 2-month-old and 6-month-old B-Cyb5r3 mice vs. RIP-Cre^+^ controls. All data are presented as means ± SEM. *p < 0.05, **p < 0.01, ***p < 0.001 by Student’s t-test, (n=3 per group).

**Supplementary Figure 5. Forest plot of disease associations with rs5758837.** The PheWAS graphic displays associations for a SNP in the *CYB5R3* promoter, rs5758837, across all phenotypes included in the Type 2 Diabetes Knowledge Portal, or across UK Biobank phenotypes [25].

**Supplementary Table 1.** List of Antibodies Used

| Antigen | Species | Dilution | Product Number | Company |
| --- | --- | --- | --- | --- |
| Insulin | Guinea Pig | 1:2000 | A0564 | DAKO |
| Aldh1a3 | Rabbit | 1:100 | NBP2-15339 | Novus |
| Cyb5r3 | Rabbit | 1:100 | 10894-1-AP | Proteintech |
| Proglucagon | Rabbit | 1:500 | #8233 | Cell Signaling |
| Somatostatin | Goat | 1:1000 | SC-7819 | Santa Cruz |
| Pancreatic Polypeptide | Goat | 1:500 | AB77192 | Abcam |
| Glut2 | Rabbit | 1:200 | AB54460 | Abcam |
| MafA | Rabbit | 1:200 | IHC00352 | Bethyl |
| Pdx1 | Goat | 1:100 | AF2419 | R&D |
| FoxO1 | Rabbit | 1:100 | CST 2880S | Cell Signaling |
| Neurogenin3 | Rabbit | 1:100 | AB2011 | BCBC |

## References

1. Accili, D., Talchai, S.C., Kim-Muller, J.Y., Cinti, F., Ishida, E., Ordelheide, A.M., et al., 2016. When beta-cells fail: lessons from dedifferentiation. Diabetes, obesity & metabolism 18 Suppl 1:117–122.

2. Halban, P.A., Polonsky, K.S., Bowden, D.W., Hawkins, M.A., Ling, C., Mather, K.J., et al., 2014. beta-cell failure in type 2 diabetes: postulated mechanisms and prospects for prevention and treatment. Diabetes Care 37(6):1751–1758.

3. Ferrannini, E., 2010. The stunned beta cell: a brief history. Cell metabolism 11(5):349–352.

4. Holst, J.J., 2019. From the Incretin Concept and the Discovery of GLP-1 to Today’s Diabetes Therapy. Front Endocrinol (Lausanne) 10:260.

5. Home, P.D., Ahren, B., Reusch, J.E.B., Rendell, M., Weissman, P.N., Cirkel, D.T., et al., 2017. Three-year data from 5 HARMONY phase 3 clinical trials of albiglutide in type 2 diabetes mellitus: Long-term efficacy with or without rescue therapy. Diabetes Res Clin Pract 131:49–60.

6. Jones, A.G., McDonald, T.J., Shields, B.M., Hill, A.V., Hyde, C.J., Knight, B.A., et al., 2016. Markers of beta-Cell Failure Predict Poor Glycemic Response to GLP-1 Receptor Agonist Therapy in Type 2 Diabetes. Diabetes Care 39(2):250–257.

7. Talchai, C., Xuan, S., Lin, H.V., Sussel, L., Accili, D., 2012. Pancreatic beta cell dedifferentiation as a mechanism of diabetic beta cell failure. Cell 150(6):1223–1234.

8. Wang, Z., York, N.W., Nichols, C.G., Remedi, M.S., 2014. Pancreatic beta cell dedifferentiation in diabetes and redifferentiation following insulin therapy. Cell Metab 19(5):872–882.

9. Spijker, H.S., Song, H., Ellenbroek, J.H., Roefs, M.M., Engelse, M.A., Bos, E., et al., 2015. Loss of β-Cell Identity Occurs in Type 2 Diabetes and Is Associated With Islet Amyloid Deposits. - PubMed - NCBI. Diabetes 64(8):2928–2938.

10. Jonas, J.C., Sharma, A., Hasenkamp, W., Ilkova, H., Patane, G., Laybutt, R., et al., 1999. Chronic hyperglycemia triggers loss of pancreatic beta cell differentiation in an animal model of diabetes. The Journal of biological chemistry 274(20):14112–14121.

11. Guo, S., Dai, C., Guo, M., Taylor, B., Harmon, J.S., Sander, M., et al., 2013. Inactivation of specific beta cell transcription factors in type 2 diabetes. J Clin Invest 123(8):3305–3316.

12. Taylor, B.L., Liu, F.F., Sander, M., 2013. Nkx6.1 is essential for maintaining the functional state of pancreatic beta cells. Cell Rep 4(6):1262–1275.

13. Puri, S., Akiyama, H., Hebrok, M., 2013. VHL-mediated disruption of Sox9 activity compromises beta-cell identity and results in diabetes mellitus. Genes Dev 27(23):2563–2575.

14. Kitamura, Y.I., Kitamura, T., Kruse, J.P., Raum, J.C., Stein, R., Gu, W., et al., 2005. FoxO1 protects against pancreatic beta cell failure through NeuroD and MafA induction. Cell Metab 2(3):153–163.

15. Kawamori, D., Kaneto, H., Nakatani, Y., Matsuoka, T.A., Matsuhisa, M., Hori, M., et al., 2006. The forkhead transcription factor Foxo1 bridges the JNK pathway and the transcription factor PDX-1 through its intracellular translocation. J Biol Chem 281(2):1091–1098.

16. Kim-Muller, J.Y., Kim, Y.J., Fan, J., Zhao, S., Banks, A.S., Prentki, M., et al., 2016. FoxO1 Deacetylation Decreases Fatty Acid Oxidation in beta-Cells and Sustains Insulin Secretion in Diabetes. J Biol Chem 291(19):10162–10172.

17. Kim-Muller, J.Y., Zhao, S., Srivastava, S., Mugabo, Y., Noh, H.L., Kim, Y.R., et al., 2014. Metabolic inflexibility impairs insulin secretion and results in MODY-like diabetes in triple FoxO-deficient mice. Cell Metab 20(4):593–602.

18. Kim-Muller, J.Y., Fan, J., Kim, Y.J., Lee, S.A., Ishida, E., Blaner, W.S., et al., 2016. Aldehyde dehydrogenase 1a3 defines a subset of failing pancreatic beta cells in diabetic mice. Nat Commun 7:12631.

19. Siendones, E., SantaCruz-Calvo, S., Martin-Montalvo, A., Cascajo, M.V., Ariza, J., Lopez-Lluch, G., et al., 2014. Membrane-bound CYB5R3 is a common effector of nutritional and oxidative stress response through FOXO3a and Nrf2. Antioxidants & redox signaling 21(12):1708–1725.

20. Shirabe, K., Landi, M.T., Takeshita, M., Uziel, G., Fedrizzi, E., Borgese, N., 1995. A novel point mutation in a 3’ splice site of the NADH-cytochrome b5 reductase gene results in immunologically undetectable enzyme and impaired NADH-dependent ascorbate regeneration in cultured fibroblasts of a patient with type II hereditary methemoglobinemia. Am J Hum Genet 57(2):302–310.

21. Martin-Montalvo, A., Sun, Y., Diaz-Ruiz, A., Ali, A., Gutierrez, V., Palacios, H.H., et al., 2016. Cytochrome b5 reductase and the control of lipid metabolism and healthspan. NPJ Aging Mech Dis 2:16006.

22. Xie, J., Zhu, H., Larade, K., Ladoux, A., Seguritan, A., Chu, M., et al., 2004. Absence of a reductase, NCB5OR, causes insulin-deficient diabetes. Proc Natl Acad Sci U S A 101(29):10750–10755.

23. Baron, M., Veres, A., Wolock, S.L., Faust, A.L., Gaujoux, R., Vetere, A., et al., 2016. A Single-Cell Transcriptomic Map of the Human and Mouse Pancreas Reveals Inter- and Intra-cell Population Structure. Cell Syst 3(4):346–360 e344.

24. Veres, A., Faust, A.L., Bushnell, H.L., Engquist, E.N., Kenty, J.H., Harb, G., et al., 2019. Charting cellular identity during human in vitro beta-cell differentiation. Nature 569(7756):368–373.

25. http://www.type2diabetesgenetics.org/variantInfo/variantInfo/22_43049014_G_T.

26. Kuo, T., Kraakman, M.J., Damle, M., Gill, R., Lazar, M.A., Accili, D., 2019. Identification of C2CD4A as a human diabetes susceptibility gene with a role in β-cell insulin secretion. Proc Natl Acad Sci U S A in press.

27. Sheng, C., Li, F., Lin, Z., Zhang, M., Yang, P., Bu, L., et al., 2016. Reversibility of beta-Cell-Specific Transcript Factors Expression by Long-Term Caloric Restriction in db/db Mouse. J Diabetes Res 2016:6035046.

28. Ishida, E., Kim-Muller, J.Y., Accili, D., 2017. Pair Feeding, but Not Insulin, Phloridzin, or Rosiglitazone Treatment, Curtails Markers of beta-Cell Dedifferentiation in db/db Mice. Diabetes 66(8):2092–2101.

29. Xiao, X., Chen, C., Guo, P., Zhang, T., Fischbach, S., Fusco, J., et al., 2017. Forkhead Box Protein 1 (FoxO1) Inhibits Accelerated beta Cell Aging in Pancreas-specific SMAD7 Mutant Mice. J Biol Chem 292(8):3456–3465.

30. Tersey, S.A., Levasseur, E.M., Syed, F., Farb, T.B., Orr, K.S., Nelson, J.B., et al., 2018. Episodic beta-cell death and dedifferentiation during diet-induced obesity and dysglycemia in male mice. The FASEB journal: official publication of the Federation of American Societies for Experimental Biology:fj201800150RR.

31. Gong, M., Yu, Y., Liang, L., Vuralli, D., Froehler, S., Kuehnen, P., et al., 2019. HDAC4 mutations cause diabetes and induce beta-cell FoxO1 nuclear exclusion. Mol Genet Genomic Med 7(5):e602.

32. Nakae, J., Kitamura, T., Kitamura, Y., Biggs, W.H., 3rd, Arden, K.C., Accili, D., 2003. The forkhead transcription factor Foxo1 regulates adipocyte differentiation. Dev Cell 4(1):119–129.

33. Whyte, W.A., Orlando, D.A., Hnisz, D., Abraham, B.J., Lin, C.Y., Kagey, M.H., et al., 2013. Master transcription factors and mediator establish super-enhancers at key cell identity genes. Cell 153(2):307–319.

34. Wollheim, C.B., Maechler, P., 2002. Beta-cell mitochondria and insulin secretion: messenger role of nucleotides and metabolites. Diabetes 51 Suppl 1:S37–42.

35. St-Pierre, J., Buckingham, J.A., Roebuck, S.J., Brand, M.D., 2002. Topology of superoxide production from different sites in the mitochondrial electron transport chain. J Biol Chem 277(47):44784–44790.

36. Laybutt, D.R., Sharma, A., Sgroi, D.C., Gaudet, J., Bonner-Weir, S., Weir, G.C., 2002. Genetic regulation of metabolic pathways in beta-cells disrupted by hyperglycemia. J Biol Chem 277(13):10912–10921.

37. Talchai, C., Xuan, S., Kitamura, T., DePinho, R.A., Accili, D., 2012. Generation of functional insulin-producing cells in the gut by Foxo1 ablation. Nature genetics 44(4):406–412, S401.

38. Accili, D., 2018. Insulin Action Research and the Future of Diabetes Treatment: The 2017 Banting Medal for Scientific Achievement Lecture. Diabetes 67(9):1701–1709.

39. Cheng, Z., Guo, S., Copps, K., Dong, X., Kollipara, R., Rodgers, J.T., et al., 2009. Foxo1 integrates insulin signaling with mitochondrial function in the liver. Nat Med 15(11):1307–1311.

40. Brand, M.D., 2016. Mitochondrial generation of superoxide and hydrogen peroxide as the source of mitochondrial redox signaling. Free Radic Biol Med.

41. Vukkadapu, S.S., Belli, J.M., Ishii, K., Jegga, A.G., Hutton, J.J., Aronow, B.J., et al., 2005. Dynamic interaction between T cell-mediated beta-cell damage and beta-cell repair in the run up to autoimmune diabetes of the NOD mouse. Physiol Genomics 21(2):201–211.

42. Ahmed, M., Muhammed, S.J., Kessler, B., Salehi, A., 2010. Mitochondrial proteome analysis reveals altered expression of voltage dependent anion channels in pancreatic beta-cells exposed to high glucose. Islets 2(5):283–292.

43. Taneera, J., Lang, S., Sharma, A., Fadista, J., Zhou, Y., Ahlqvist, E., et al., 2012. A systems genetics approach identifies genes and pathways for type 2 diabetes in human islets. Cell Metab 16(1):122–134.

44. Solimena, M., Schulte, A.M., Marselli, L., Ehehalt, F., Richter, D., Kleeberg, M., et al., 2018. Systems biology of the IMIDIA biobank from organ donors and pancreatectomised patients defines a novel transcriptomic signature of islets from individuals with type 2 diabetes. Diabetologia 61(3):641–657.

45. Xin, Y., Kim, J., Okamoto, H., Ni, M., Wei, Y., Adler, C., et al., 2016. RNA Sequencing of Single Human Islet Cells Reveals Type 2 Diabetes Genes. Cell Metab.

46. Anello, M., Lupi, R., Spampinato, D., Piro, S., Masini, M., Boggi, U., et al., 2005. Functional and morphological alterations of mitochondria in pancreatic beta cells from type 2 diabetic patients. Diabetologia 48(2):282–289.

47. Bindokas, V.P., Kuznetsov, A., Sreenan, S., Polonsky, K.S., Roe, M.W., Philipson, L.H., 2003. Visualizing superoxide production in normal and diabetic rat islets of Langerhans. J Biol Chem 278(11):9796–9801.

48. Lowell, B.B., Shulman, G.I., 2005. Mitochondrial dysfunction and type 2 diabetes. Science 307(5708):384–387.

49. Gauthier, B.R., Wiederkehr, A., Baquie, M., Dai, C., Powers, A.C., Kerr-Conte, J., et al., 2009. PDX1 deficiency causes mitochondrial dysfunction and defective insulin secretion through TFAM suppression. Cell Metab 10(2):110–118.

50. Maechler, P., Li, N., Casimir, M., Vetterli, L., Frigerio, F., Brun, T., 2010. Role of mitochondria in beta-cell function and dysfunction. Advances in experimental medicine and biology 654:193–216.

51. Lenzen, S., Drinkgern, J., Tiedge, M., 1996. Low antioxidant enzyme gene expression in pancreatic islets compared with various other mouse tissues. Free Radic Biol Med 20(3):463–466.

52. Elahian, F., Sepehrizadeh, Z., Moghimi, B., Mirzaei, S.A., 2014. Human cytochrome b5 reductase: structure, function, and potential applications. Crit Rev Biotechnol 34(2):134–143.

53. Jimenez-Hidalgo, M., Santos-Ocana, C., Padilla, S., Villalba, J.M., Lopez-Lluch, G., Martin-Montalvo, A., et al., 2009. NQR1 controls lifespan by regulating the promotion of respiratory metabolism in yeast. Aging Cell 8(2):140–151.

54. Busch, A.K., Gurisik, E., Cordery, D.V., Sudlow, M., Denyer, G.S., Laybutt, D.R., et al., 2005. Increased fatty acid desaturation and enhanced expression of stearoyl coenzyme A desaturase protects pancreatic beta-cells from lipoapoptosis. Diabetes 54(10):2917–2924.

55. Abebe, T., Mahadevan, J., Bogachus, L., Hahn, S., Black, M., Oseid, E., et al., 2017. Nrf2/antioxidant pathway mediates beta cell self-repair after damage by high-fat diet-induced oxidative stress. JCI Insight 2(24).

56. Kitamura, T., Kitamura, Y.I., Kobayashi, M., Kikuchi, O., Sasaki, T., Depinho, R.A., et al., 2009. Regulation of pancreatic juxtaductal endocrine cell formation by FoxO1. Mol Cell Biol 29(16):4417–4430.

57. Tsuchiya, K., Tanaka, J., Shuiqing, Y., Welch, C.L., DePinho, R.A., Tabas, I., et al., 2012. FoxOs integrate pleiotropic actions of insulin in vascular endothelium to protect mice from atherosclerosis. Cell Metab 15(3):372–381.

58. Lin, H.V., Ren, H., Samuel, V.T., Lee, H.Y., Lu, T.Y., Shulman, G.I., et al., 2011. Diabetes in mice with selective impairment of insulin action in Glut4-expressing tissues. Diabetes 60(3):700–709.

59. Postic, C., Shiota, M., Niswender, K.D., Jetton, T.L., Chen, Y., Moates, J.M., et al., 1999. Dual roles for glucokinase in glucose homeostasis as determined by liver and pancreatic beta cell-specific gene knock-outs using Cre recombinase. J Biol Chem 274(1):305–315.

60. Nakae, J., Kitamura, T., Silver, D.L., Accili, D., 2001. The forkhead transcription factor Foxo1 (Fkhr) confers insulin sensitivity onto glucose-6-phosphatase expression. J Clin Invest 108(9):1359–1367.

61. Son, J., Shen, S.S., Margueron, R., Reinberg, D., 2013. Nucleosome-binding activities within JARID2 and EZH1 regulate the function of PRC2 on chromatin. Genes Dev 27(24):2663–2677.

62. Kuo, T., Damle, M., Gonzalez, B.J., Egli, D., Lazar, M.A., Accili, D., 2019. Induction of alpha cell-restricted Gc in dedifferentiating beta cells contributes to stress-induced beta-cell dysfunction. JCI Insight 5.

63. Creyghton, M.P., Cheng, A.W., Welstead, G.G., Kooistra, T., Carey, B.W., Steine, E.J., et al., 2010. Histone H3K27ac separates active from poised enhancers and predicts developmental state. Proc Natl Acad Sci U S A 107(50):21931–21936.

64. Kitamura, T., Kido, Y., Nef, S., Merenmies, J., Parada, L.F., Accili, D., 2001. Preserved pancreatic beta-cell development and function in mice lacking the insulin receptor-related receptor. Mol Cell Biol 21(16):5624–5630.

65. Nakae, J., Biggs, W.H., 3rd, Kitamura, T., Cavenee, W.K., Wright, C.V., Arden, K.C., et al., 2002. Regulation of insulin action and pancreatic beta-cell function by mutated alleles of the gene encoding forkhead transcription factor Foxo1. Nature genetics 32(2):245–253.

66. Spinazzi, M., Casarin, A., Pertegato, V., Salviati, L., Angelini, C., 2012. Assessment of mitochondrial respiratory chain enzymatic activities on tissues and cultured cells. Nat Protoc 7(6):1235–1246.

67. Haeusler, R.A., Kaestner, K.H., Accili, D., 2010. FoxOs function synergistically to promote glucose production. J Biol Chem 285(46):35245–35248.

68. Plum, L., Lin, H.V., Dutia, R., Tanaka, J., Aizawa, K.S., Matsumoto, M., et al., 2009. The obesity susceptibility gene Cpe links FoxO1 signaling in hypothalamic pro-opiomelanocortin neurons with regulation of food intake. Nat Med 15(10):1195–1201.

