## Supplemental Figures for "Cyb5r3 links FoxO1-dependent mitochondrial dysfunction with β-cell failure"

### Slide 1
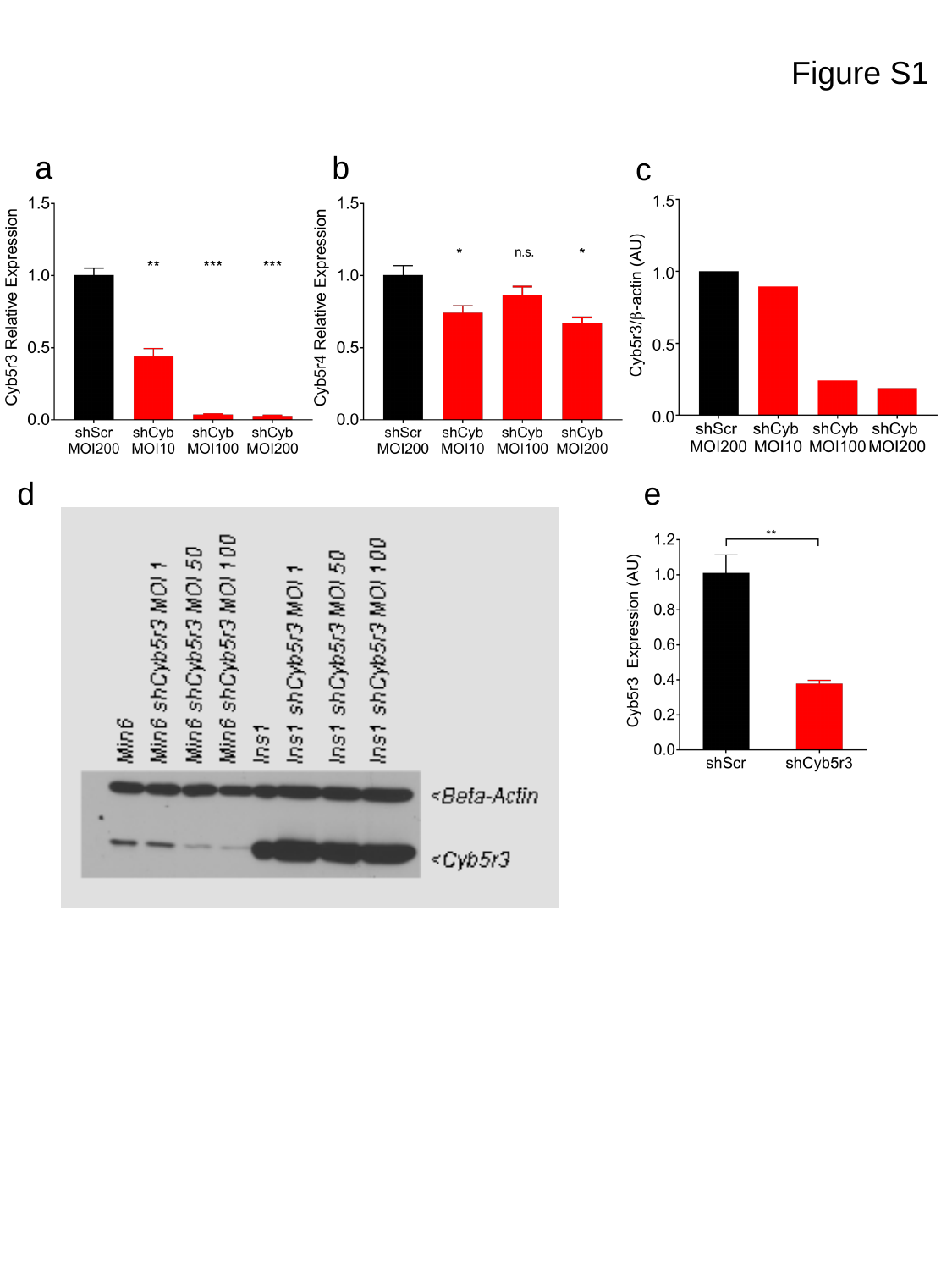

Figure S1
a
b
d
c
e

### Slide 2
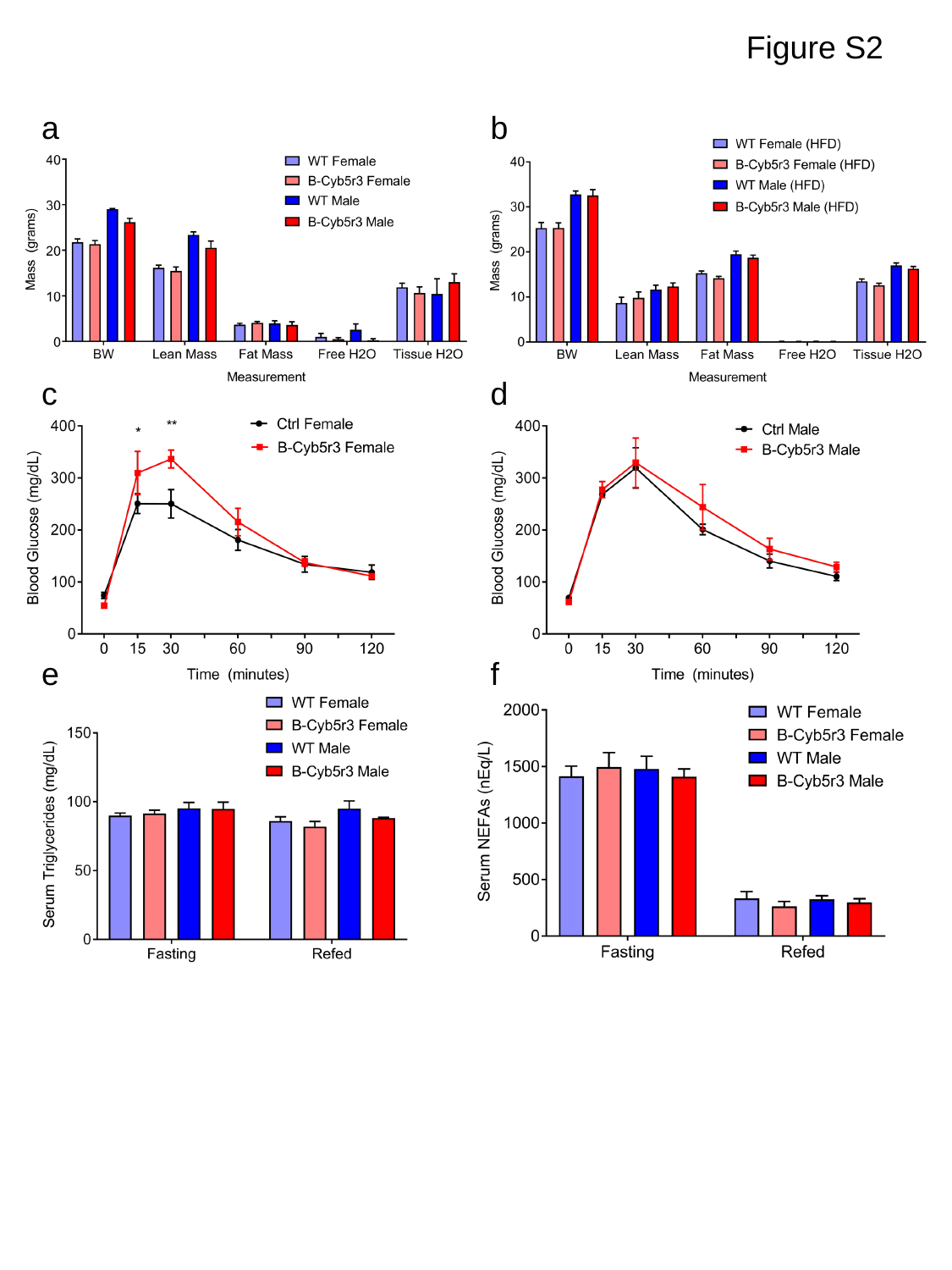

Figure S2
a
b
c
d
e
f

### Slide 3
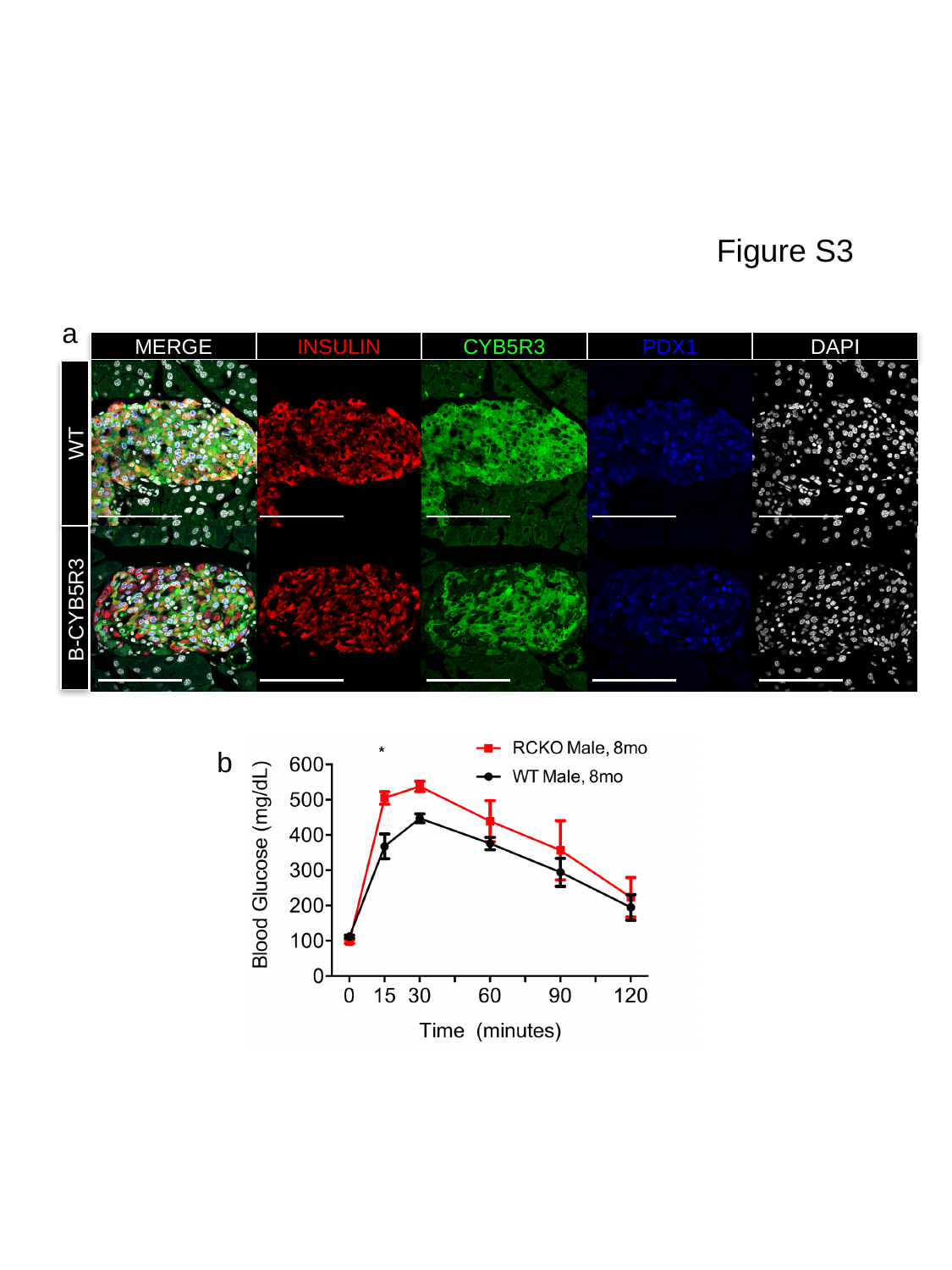

Figure S3
a
MERGE
INSULIN
CYB5R3
PDX1
DAPI
WT
B-CYB5R3
b

### Slide 4
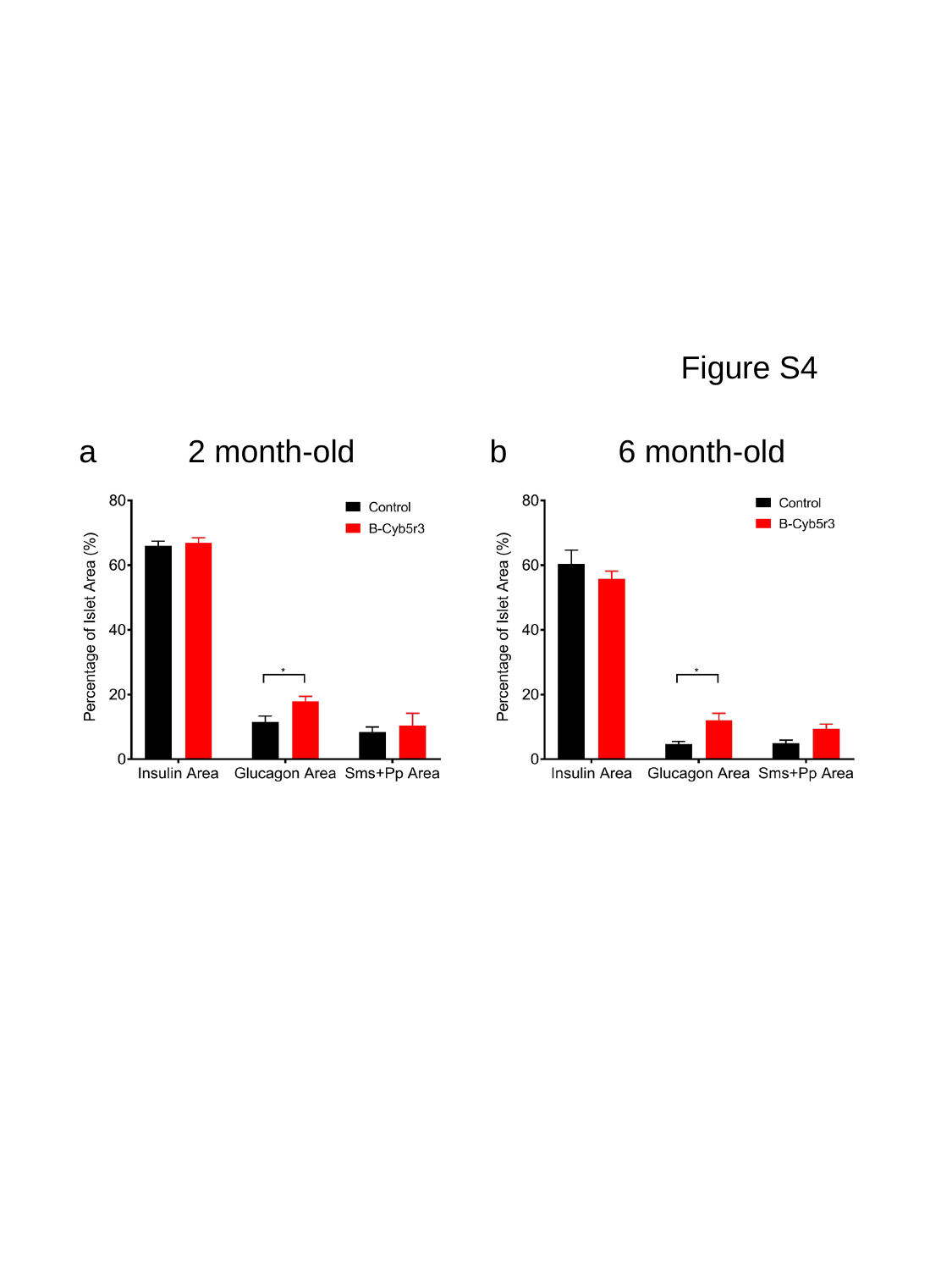

Figure S4
a
b
2 month-old
6 month-old

### Slide 5
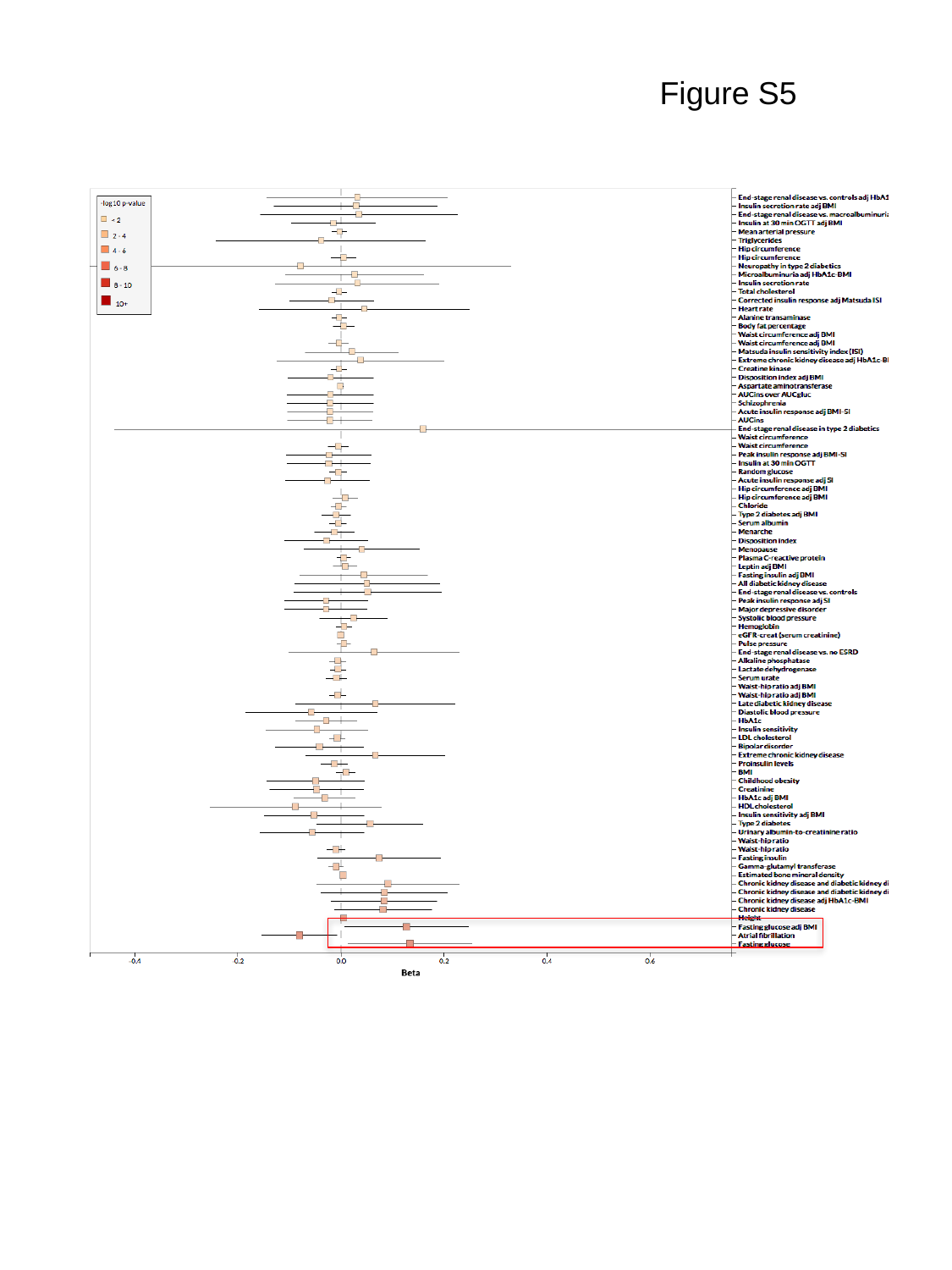

Figure S5
